# Improving Emotion Classification by Combining fNIRS-Derived Hemodynamic Responses with Peripheral Physiological Signals

**DOI:** 10.64898/2026.04.02.714099

**Authors:** Shigeyuki Ikeda, Sho Tsukawaki, Takayuki Nozawa

## Abstract

We investigated whether multimodal sensing that combines functional near-infrared spectroscopy (fNIRS) with peripheral physiological signals can improve subject-independent classification of arousal and valence, the fundamental affective dimensions in Russell’s circumplex model. We developed Japanese emotion-inducing music-video stimuli (60 seconds each) and recorded subjects’ central nervous system activity using fNIRS, alongside peripheral physiological measures, specifically electrodermal activity (EDA) and photoplethysmography (PPG), during video viewing. To prioritize reproducibility and methodological transparency, we extracted simple, easily computed features from each modality and performed binary (high vs. low) classification separately for arousal and valence using a support vector machine. The combination of fNIRS and EDA yielded the highest performance, with a macro-averaged F1 score of 0.73 for arousal and 0.64 for valence. These findings underscore the utility of integrating fNIRS with peripheral physiological signals for subject-independent emotion classification.

## I. INTRODUCTION

Accurately and quantitatively inferring human affect is a core enabling capability for affective computing, with direct implications for adaptive user interfaces and applications that support human well-being. One of the most established approaches to assessing affect is self-report using psychometric instruments. Representative examples include the Self-Assessment Manikin (SAM) [1] and the Positive and Negative Affect Schedule (PANAS) [2]. Although questionnaires can capture subjective experience directly and therefore remain a strong reference standard, repeated self-reporting imposes non-negligible burden and is often impractical in settings that require continuous or frequent assessment.

To mitigate this limitation, prior work has investigated automated affect inference by combining external sensing of human behavioral and physiological responses with machine learning [3]. Common sensing targets include facial expressions, speech, body movements, and physiological signals. Facial, vocal, and behavioral cues provide informative outward manifestations of affect; however, they may be attenuated or absent when emotions are internally experienced but not overtly expressed. In contrast, physiological sensing can provide a more direct window into internal affective processes.

Among physiological modalities, electroencephalography (EEG) has been extensively studied for affect recognition (see [4] for a review; representative studies include [5], [6], [7], [8], [9]). Because EEG measures central nervous system activity more directly than peripheral signals, it can be sensitive to rapid affective changes [10], [11]. EEG systems can also be relatively portable, and multimodal combinations of EEG with peripheral measures have been reported to improve recognition performance (see [12] for a review). Nevertheless, EEG recordings are highly susceptible to artifacts from body motion, ocular activity (e.g., blinks), and electromagnetic interference from the environment. These factors present substantial challenges for robust affective computing in everyday settings.

Functional near-infrared spectroscopy (fNIRS) provides an alternative approach to measuring brain activity. fNIRS quantifies changes in oxygenated and deoxygenated hemoglobin concentrations associated with cerebral hemodynamics. fNIRS signals reflect a slower hemodynamic response that is indirectly coupled to neuronal activity, resulting in lower temporal resolution than electrophysiological measures [13]. On the other hand, compared with EEG, fNIRS is generally more tolerant to certain motion- and blink-related artifacts and is not affected by electromagnetic noise, which makes it attractive for measurements in more naturalistic environments [14], [15]. In fact, multiple studies have demonstrated successful affect recognition using fNIRS alone [16], [17], [18], [19], and subject-independent models for arousal and valence have also been reported [20]. More recently, multimodal affect recognition combining fNIRS and EEG has been explored [21], [22]. In contrast, it remains less clear to what extent combining fNIRS with peripheral autonomic measures yields consistent benefits for affect recognition, particularly under subject-independent evaluation.

A related study reported that combining fNIRS with electrodermal activity (EDA) enables coarse estimation of arousal levels. However, it remains unclear whether this multimodal approach yields a measurable improvement over unimodal fNIRS or EDA, and whether a subject-independent estimation model can be learned [23]. Another closely related recent study investigated multimodal affect recognition using multiple sensors, including electrocardiogram, EDA, respiration, skin temperature, fNIRS, and EEG, to classify five emotion categories [24]. This work reported high performance for subject-dependent models, while highlighting the difficulty of learning subject-independent models and leaving open the question of which peripheral modalities are most complementary to fNIRS. In response to these identified gaps, the current study examines multimodal emotion classification utilizing fNIRS to assess central nervous system activity, alongside two commonly employed peripheral measures of autonomic function—EDA and photoplethysmography (PPG). The investigation is centered on the core affective dimensions of arousal and valence as delineated in Russell’s circumplex model [25]. Specifically, we evaluate whether combining EDA and/or PPG with fNIRS improves classification performance relative to fNIRS alone, and whether subject-independent (cross-subject) emotion classification can be achieved using fNIRS-centered multimodal sensing. To prioritize reproducibility and transparency, we restrict the input representation to commonly used, simple features (e.g., mean and standard deviation) and features that can be computed using widely adopted toolboxes, and we employ classical machine-learning classifiers rather than highly parameterized approaches such as deep learning.

## II. Material and methods

### A. Subjects

The subjects included 35 Japanese, healthy, right-handed, university or postgraduate students (24 men and 11 women, mean age: 20.7 ± 1.26 years; mean ± SD). All subjects either had normal vision without corrective lenses or wore contact lenses during the experiment. We obtained written informed consent from all subjects.

### B. Stimuli Selection

We assembled an initial pool of 40 Japanese popular music videos from YouTube. From each video, we extracted a 60-s segment expected to elicit affective responses (e.g., the chorus), resulting in 40 stimuli of equal duration. The stimuli were shown in Supplementary Table S1.

To obtain subjective affective labels, 10 subjects (out of the total sample of 35) watched all 40 stimuli and rated their experienced arousal and valence using the 9-point SAM scales. However, one subject did not complete the experiment due to a malfunction in stimulus-presentation. Based on the ratings of the remaining 9 subjects, we selected 12 stimuli that elicited comparatively strong and reliable affective responses.

Stimulus selection proceeded as follows. First, SAM ratings were mean-centered by subtracting 5 from the original 1–9 scores. Next, for each stimulus, we computed the mean and standard deviation of the centered ratings across the 9 subjects and defined a normalized emotional score as the ratio of the mean to the standard deviation (mean/SD) [9]. We then ranked the 40 stimuli according to the normalized emotional score and selected 12 stimuli with the highest (or lowest) scores (Fig. 1).

**Fig. 1.**
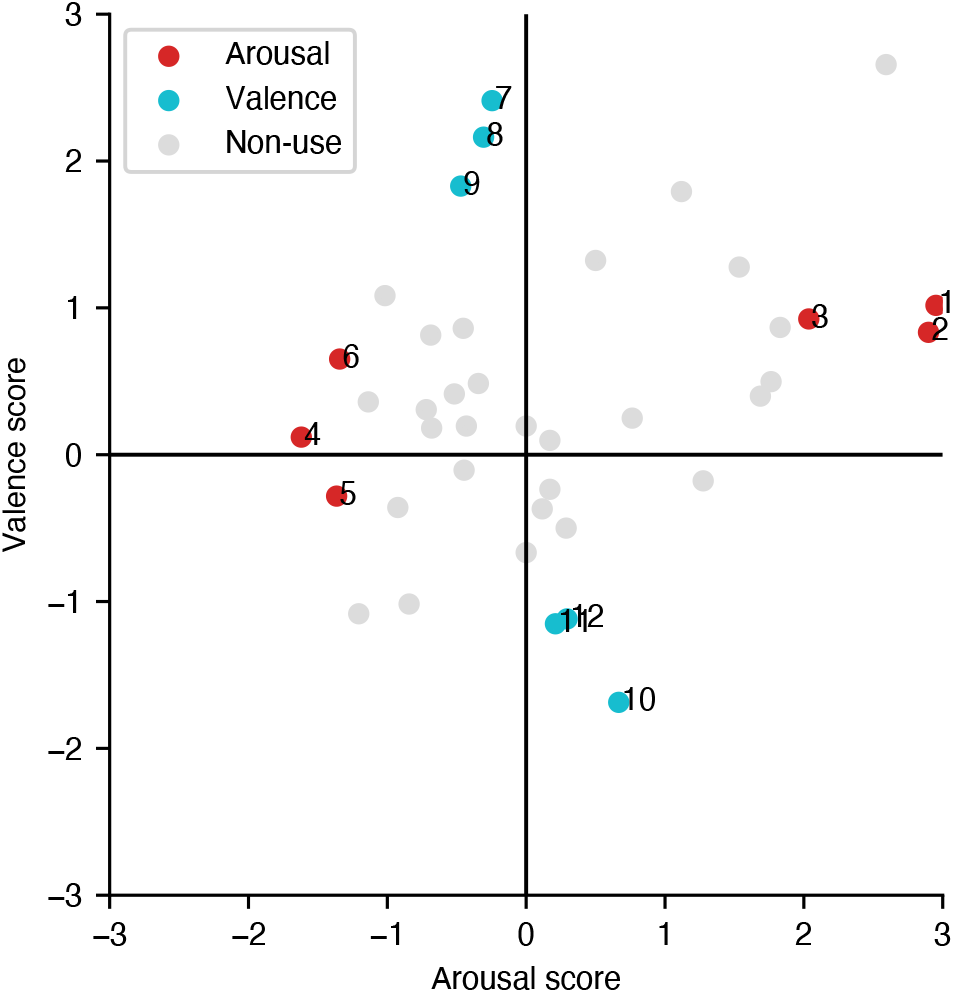
Scatter plot of the 40 candidate music-video stimuli based on the normalized emotional scores. Each point represents one 60-s music-video segment positioned in the two-dimensional space defined by the normalized emotional scores for arousal and valence. Each Arabic numeral corresponds to the number of each video shown in Supplementary Table S1. Red points indicate stimuli selected to elicit relatively high or low arousal. Cyan points indicate stimuli selected to elicit relatively high or low valence. Light gray points denote candidate stimuli that were not selected for the main experiment. A candidate located in the upper-right region of the plot was excluded because they elicited simultaneously strong arousal and strong valence, which could confound dimension-specific stimulus selection.

The final stimulus set comprised three videos in each of four affective categories: high/low arousal and high/low valence. Additional details of the 12 selected stimuli are provided in Supplementary Table S1.

### C. fNIRS Measurement

Brain activity was recorded using a 34-channel wearable fNIRS system (WOT-HS; Hitachi High-Technologies Corporation, Japan). The system comprises 12 light sources and 12 detectors, arranged with a source–detector separation of 32 mm. Dual-wavelength light-emitting diodes were used at 730 nm and 850 nm. The measurement targeted changes in oxy- and deoxy-hemoglobin concentrations. Signals were sampled at 10 Hz.

The 34 measurement channels covered portions of the frontal and temporal cortices. To facilitate region-level analyses, we determined the location of each channel in standard space (Montreal Neurological Institute; MNI coordinates) and partitioned the 34 channels into five anatomical regions based on the automated anatomical labeling [26]: right temporal area, right lateral prefrontal area, medial prefrontal area, left lateral prefrontal area, and left temporal area (Fig. 2).

**Fig. 2.**
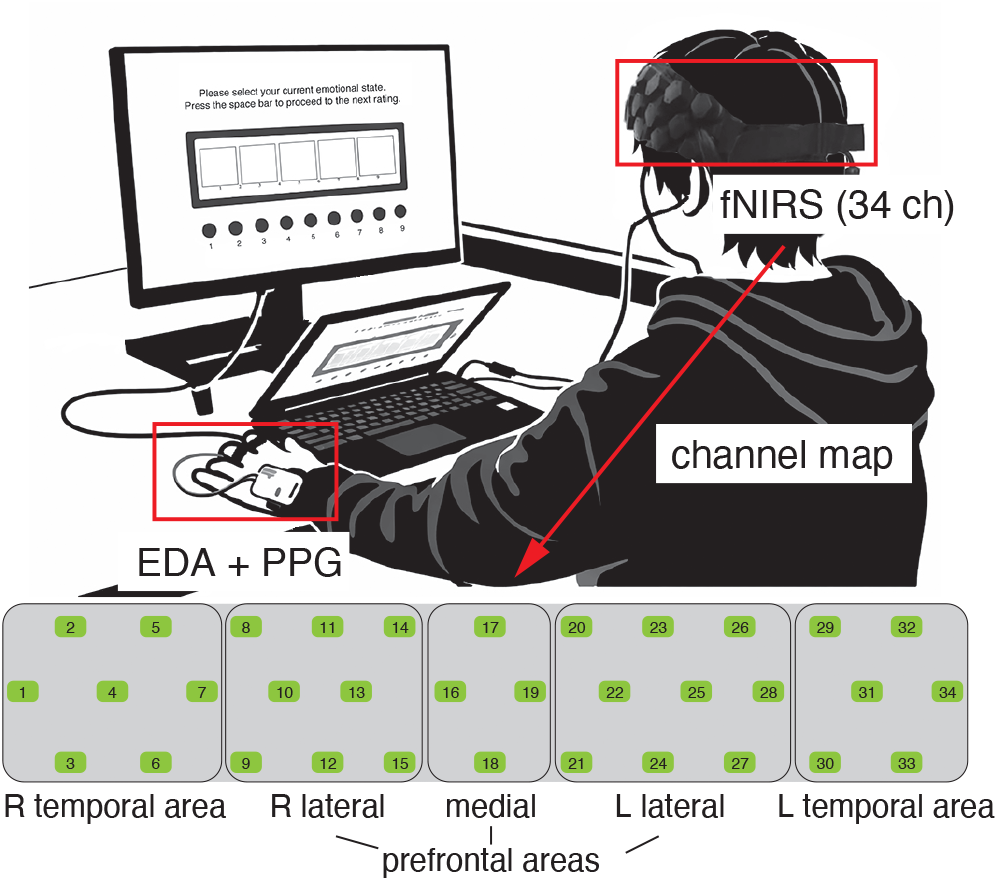
A subject wearing the fNIRS, EDA, PPG sensors and brain regions covered by the fNIRS probes.

### D. EDA and PPG Measurements

EDA and PPG signals were recorded using a Shimmer3 GSR+ Unit (Shimmer, Dublin) equipped with the GSR+ module for EDA and an optical pulse sensor for PPG. For EDA acquisition, snap-connector Ag/AgCl electrodes were placed on the ventral surfaces of the left index and middle fingers. PPG signals were measured from the fingertip of the left index finger using the optical pulse sensor. Both EDA and PPG signals were sampled at 51.2 Hz and were acquired simultaneously within the same device, ensuring hardware-level synchronization between modalities. Data were recorded using Shimmer Consensys (v1.6.0).

### E. Experimental Protocol

The objective of this experiment was to present emotion-eliciting videos to subjects wearing wired earphones and to record their central and peripheral physiological activity using fNIRS, EDA, and PPG sensors during video viewing.

A total of 35 subjects were enrolled. Ten subjects viewed 40 music-video stimuli (60 s each), during which fNIRS, EDA, and PPG signals were recorded. The experiment comprised four sessions, each including 10 videos. Each session began with a 20-s resting baseline. After each video, subjects rated their subjective arousal and valence using the 9-point SAM scales, followed by a 20-s resting interval. We utilized the PsychoPy2 software (v2023.2.3) to develop the experimental stimuli involving music videos and the SAM scales [27].

As described in the Stimulus Selection section, the affective ratings obtained from nine of the ten subjects were used to select 12 videos that elicited stronger and more consistent affective responses.

The remaining 25 subjects completed a single-session experiment in which they viewed the 12 selected videos while fNIRS, EDA, and PPG signals were recorded. This session began with a 20-s resting baseline. After each video, subjects rated arousal and valence on the 9-point SAM scales, followed by a 20-s resting interval.

### F. Data Preprocessing and Feature Extraction

Data analyses were conducted using MATLAB R2025b and Python 3.9.19.

Data from five of the 35 subjects were excluded from analysis due to experimental or recording failures.

Specifically, one subject did not complete the experiment because of a stimulus-presentation malfunction, as noted in the Stimulus Selection section. Two subjects were excluded due to fNIRS recording issues, and the remaining two subjects were excluded due to Shimmer (EDA/PPG) recording failures. As a result, data from the remaining 30 subjects were used for subsequent analyses.

*1)fNIRS:* fNIRS preprocessing was performed on a per-subject basis. First, channels exhibiting excessive noise were identified by visual inspection and excluded from subsequent analyses. To remove motion-related artifacts, we applied a wavelet-based motion artifact correction method with the interquartile-range tuning parameter set to *iqr* = 1.5 [28]. A comprehensive prior review has reported that wavelet-based correction with *iqr* values between 1.2 and 1.5 provides among the best performance for suppressing spike-like artifacts in fNIRS recordings [29]. Wavelet-based correction was implemented using Homer3 (v1.87.0) [30].

After motion artifact correction, signals were low-pass filtered at 0.5 Hz. We then removed slow trends by fitting and subtracting a third-order polynomial from each channel.

Finally, signals from each channel were standardized using z-score normalization.

The 34 channels were averaged into five brain regions: medial prefrontal, bilateral lateral prefrontal, and bilateral temporal regions, by computing the mean signals from channels corresponding to each brain region.

From the fNIRS signals recorded during each 60-s video-viewing interval, we computed three basic types of features: the mean, standard deviation, and the temporal slope over the interval (i.e., the linear trend from stimulus onset to offset). Features were extracted separately from the oxygenated hemoglobin and deoxygenated hemoglobin signals within each of the five predefined cortical regions. Consequently, the total number of fNIRS features was 5 regions × 2 hemoglobin signals × 3 features = 30 per video.

*2) EDA:* Prior to feature extraction, the EDA signal was upsampled to 256 Hz (= 51.2 Hz × 5) because subsequent processing with NeuroKit2 (v0.2.10) [31] requires the sampling rate to be specified as an integer-valued frequency.

As one feature for emotion classification, we computed the autocorrelation of the EDA signal at a 4-s time lag, which corresponds to the default setting in NeuroKit2. We then decomposed the EDA signal into its tonic and phasic components using NeuroKit2. From each component, we extracted three distributional features: standard deviation, skewness, and kurtosis. In addition, from the phasic component, we computed the number of peaks and the mean peak amplitude, both using NeuroKit2’s peak-detection routines.

In total, nine EDA features were derived per video-viewing interval and used for subsequent emotion classification analyses.

*3) PPG:* Similar to the EDA preprocessing, the PPG signal recorded during each video-viewing interval was upsampled to 256 Hz (i.e., 51.2 Hz × 5) prior to feature extraction. We then extracted 27 features from the PPG signal using NeuroKit2. These features comprised 19 time-domain and 8 frequency-domain heart-rate-variability metrics. All features were computed using NeuroKit2’s built-in function (ppg_analyze). Detailed definitions of the extracted features are provided in Supplementary Table S2.

### G. Emotion Classification Procedures

Classification procedures were performed using scikit-learn (v1.2.2).

For each subject, we obtained 12 samples in total: six samples for arousal (high/low) and six samples for valence (high/low). First, for each subject and for each target dimension (arousal or valence), we z-score normalized each feature across the corresponding six samples. Next, the standardized samples were concatenated across all subjects (*N* = 30), yielding 180 samples per dimension for classification.

We then performed two-class classification (high vs. low) separately for arousal and valence using grouped 5-fold cross-validation, where the group corresponded to subject identity. This grouping ensured that samples from the same subject were never split between the training and test sets.

In each cross-validation fold, we trained a support vector machine (SVM) classifier with a radial basis function kernel using the training set, without performing hyperparameter tuning. SVMs are most commonly used for emotion recognition [32].

Classification performance was evaluated using macro-averaged F1, macro-averaged precision, macro-averaged recall, and accuracy. To assess the variability of performance, we repeated the entire cross-validation procedure 20 times with randomized train–test fold assignments. The median was computed across the 20 classification performance scores.

Classification was performed using feature sets derived from a single modality (fNIRS, EDA, or PPG) as well as from multimodal combinations of these modalities (i.e., fNIRS + EDA, fNIRS + PPG, EDA + PPG, and fNIRS + EDA + PPG).

### H. Permutation Test of Classification Performance

To evaluate the statistical significance of classification performance, we conducted a permutation test. Specifically, to preserve the within-subject data structure, the high/low labels were randomly permuted within each subject. For each permuted dataset, we performed grouped 5-fold cross-validation repeated 20 times, using the same procedure as in the main analysis (i.e., subject identity was used as the grouping variable to prevent samples from the same subject appearing in both training and test sets).

This permutation procedure was repeated 300 times to obtain a null distribution of median classification performance values. The p-value was computed as the probability that the null distribution yielded a median greater than or equal to that obtained with the original (unpermuted) labels. Classification performance was deemed statistically significant when *p* < 0.05.

### I. Statistical Tests for Comparing Feature Sets

For each feature set, we obtained 20 classification performance scores from the repeated cross-validation procedure. To assess statistically significant differences in classification performance between feature sets, we performed a two-sided Wilcoxon signed-rank test for all pairwise comparisons between feature sets. Differences were deemed statistically significant when *p* < 0.05. These procedures were applied to each performance metric: macro-averaged F1, macro-averaged precision, macro-averaged recall, and accuracy.

### J. Permutation Feature Importance

To quantify the contribution of individual features to emotion classification, we computed permutation feature importance (hereafter, permutation importance) using scikit-learn’s permutation_importance function. Specifically, after training the classifier, we evaluated its performance on the held-out test set and then re-evaluated performance on a corrupted version of the same test set in which the values of a single feature were randomly permuted across samples. The permutation importance for that feature was defined as the decrease in performance from the original to the permuted test set; larger decreases indicate a greater contribution of the feature to classification.

Permutation importance was computed within the grouped 5-fold cross-validation framework repeated 20 times, using the same train–test splitting procedure as in the main analysis. The median was determined across the 20 permutation importance values of individual features.

To assess statistical significance of each permutation importance, we performed a permutation test, using the same procedure as in the Permutation Test of Classification Performance section. Specifically, for each permuted dataset in which the high/low labels were randomly permuted within each subject, permutation importance of individual features was computed. This process results in the generation of a null distribution of each feature’s permutation importance. The p-value of each feature was computed as the probability that the null distribution yielded a median greater than or equal to that obtained with the original (unpermuted) labels. Permutation importance was considered statistically significant when *p* < 0.05.

## III. Results

### A. Subjective Emotion Ratings

To examine whether the 12 selected music-video stimuli elicited the intended affective responses across all thirty-subjects, we visualized the stimuli in a scatter plot based on the SAM ratings of arousal and valence (Fig. 3). Ratings were normalized using the same procedure described in the Stimulus Selection section. Overall, the 12 stimuli exhibited the expected affective rating pattern.

**Fig. 3.**
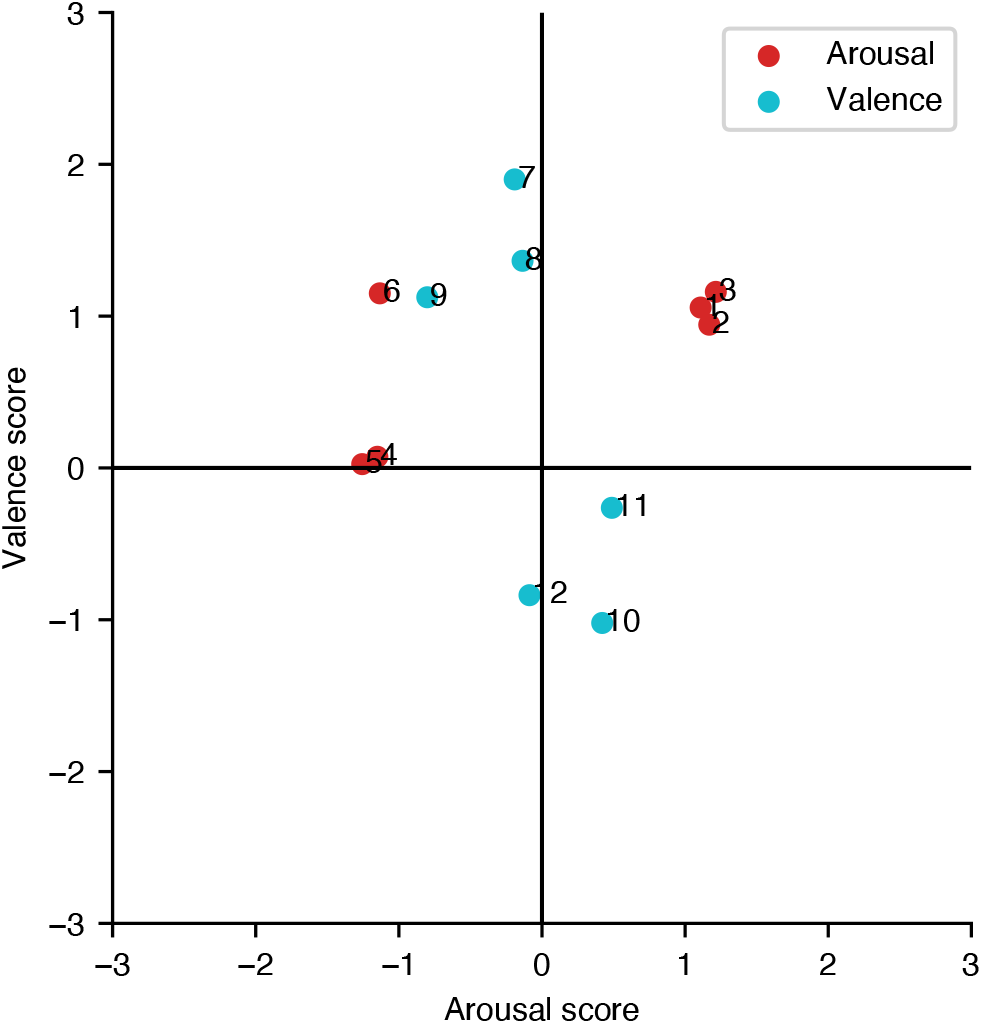
Scatter plot of the 12 music-video stimuli based on the normalized emotional scores. Each point represents one 60-s music-video segment positioned in the two-dimensional space defined by the normalized emotional scores for arousal and valence. Each Arabic numeral corresponds to the number of each video shown in Supplementary Table S1. Red points indicate stimuli selected to elicit relatively high or low arousal. Cyan points indicate stimuli selected to elicit relatively high or low valence.

### B. Emotion Classification and Comparison of Feature Sets

For arousal macro-averaged F1 scores, all feature sets achieved performance significantly above chance, with median values as follows: fNIRS = 0.65; EDA = 0.69; PPG = 0.59; fNIRS + EDA = 0.73; fNIRS + PPG = 0.69; EDA + PPG = 0.67; fNIRS + EDA + PPG = 0.72 (Fig. 4A). The fNIRS + EDA feature set yielded the highest arousal classification performance. Pairwise comparisons further showed that fNIRS + EDA outperformed all other feature sets, except the full multimodal feature set (fNIRS + EDA + PPG) (Fig. 4B).

**Fig. 4.**
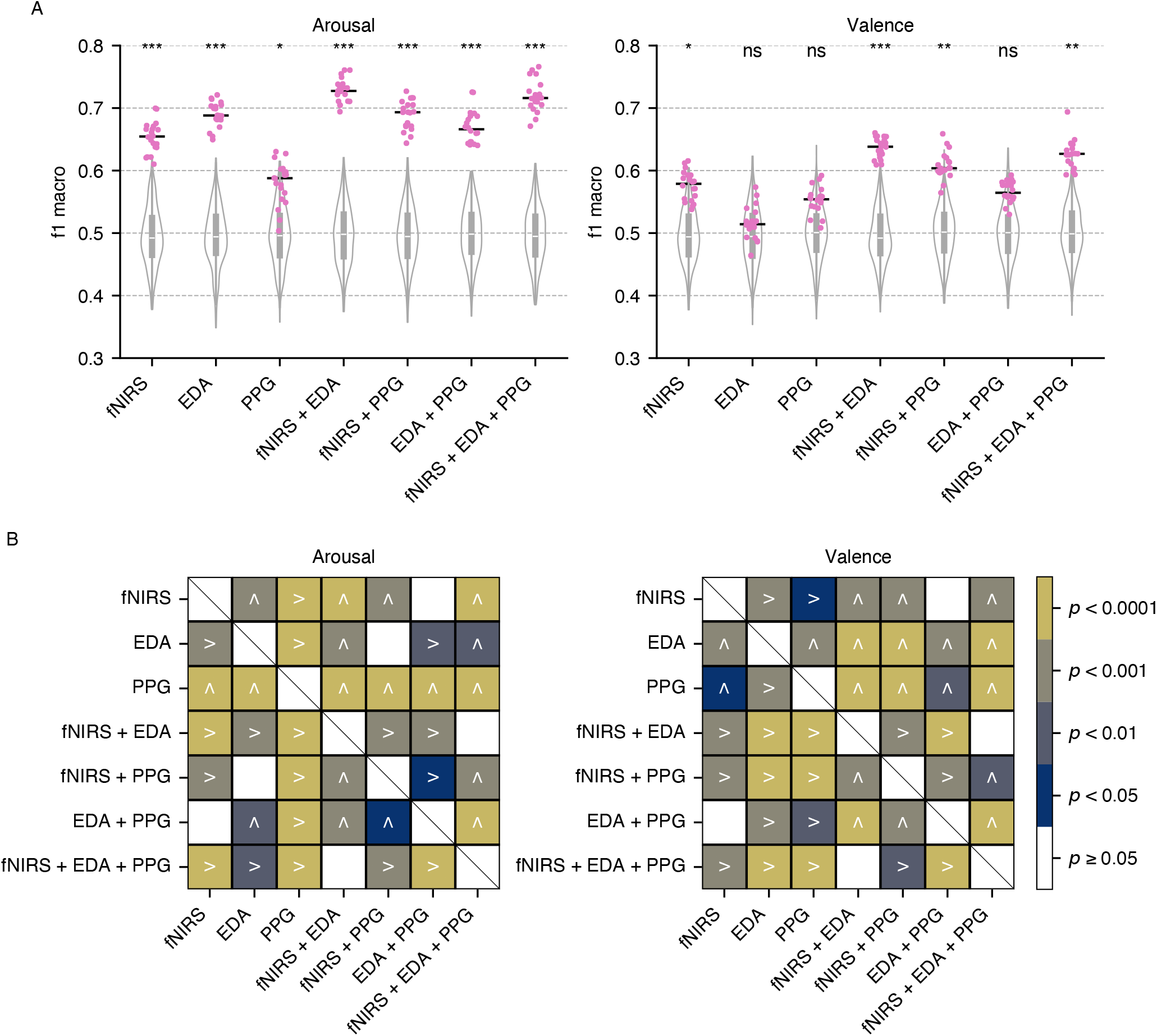
Results of emotion classification. (A) Macro-averaged F1 scores for each feature set. Pink dots denote the 20 performance estimates obtained from repeated grouped 5-fold cross-validation, and the horizontal black line indicates the median across repetitions. Violin plots show the null distributions generated by the permutation test. Asterisks indicate significance levels of the observed performance relative to the null distribution (ns: *p* ≥ 0.05; * *p* < 0.05; ** *p* < 0.01; *** *p* < 0.005). (B) Pairwise comparisons of macro-averaged F1 scores between feature sets; colors indicate significance levels from the two-sided Wilcoxon signed-rank test. The symbol “>” denotes that the feature set in the row outperformed that in the column, whereas the “∧” symbol denotes the opposite (i.e., the feature set in the column outperformed that in the row).

Valence classification proved to be somewhat more difficult than arousal classification, with the following median values of the macro-averaged F1 scores: fNIRS = 0.58; EDA = 0.51; PPG = 0.55; fNIRS + EDA = 0.64; fNIRS + PPG = 0.6; EDA + PPG = 0.56; fNIRS + EDA + PPG = 0.63. Nevertheless, the fNIRS + EDA feature set again achieved the highest performance and outperformed the other feature sets, with the exception of fNIRS + EDA + PPG.

Results for additional evaluation metrics are provided in the Supplementary Material: macro-averaged precision (Fig. S1), macro-averaged recall (Fig. S2), and accuracy (Fig. S3). The same overall pattern observed in Fig. 4 was also evident across these metrics.

### C. Permutation Importance of Each Feature

The fNIRS + EDA feature set achieved higher classification performance than the other feature sets, with the exception of the full multimodal feature set (fNIRS + EDA + PPG). Given this performance and the compactness of the fNIRS + EDA feature representation, we therefore examined permutation importance for the fNIRS + EDA model (Fig. 5).

**Fig. 5.**
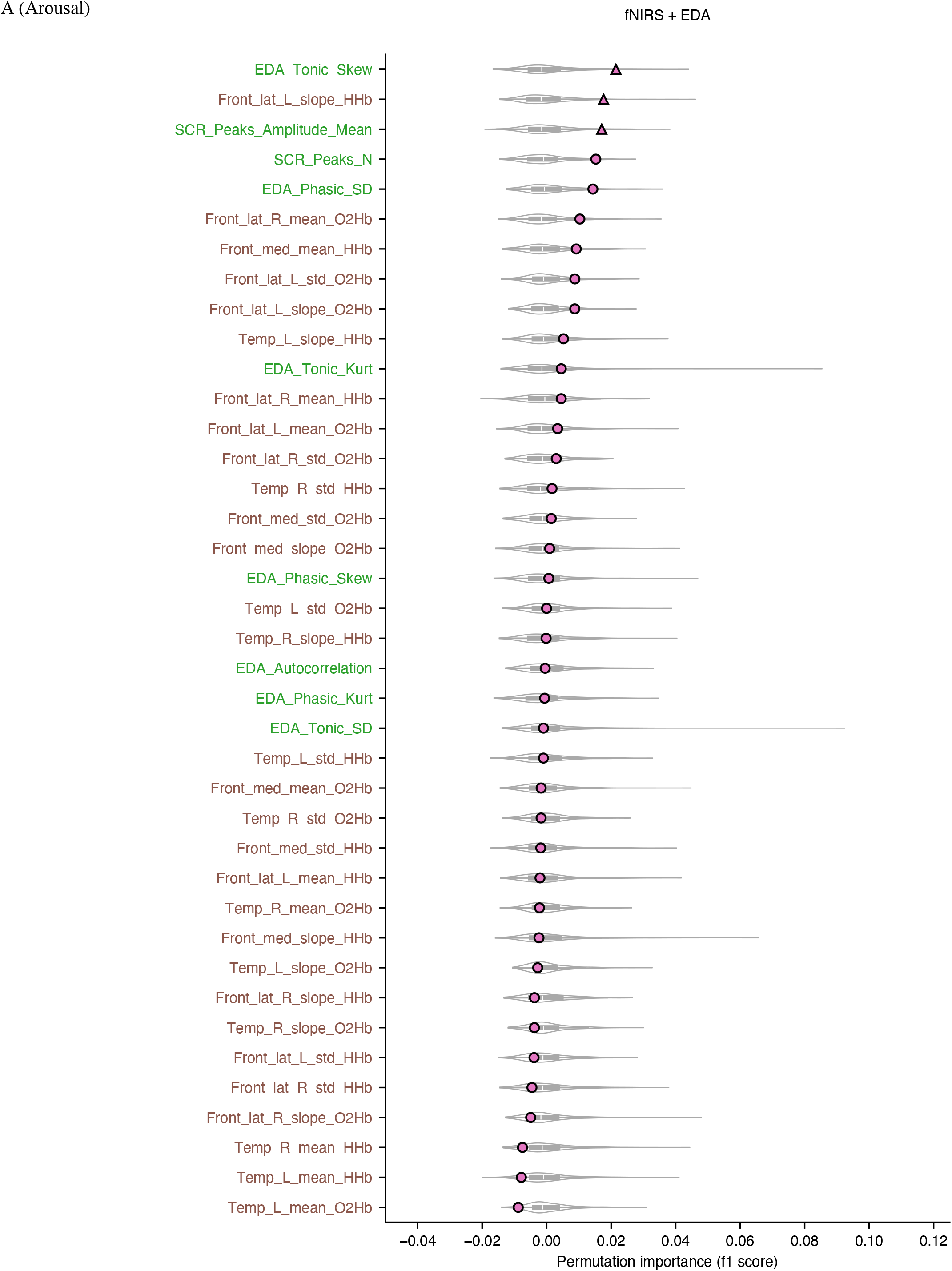
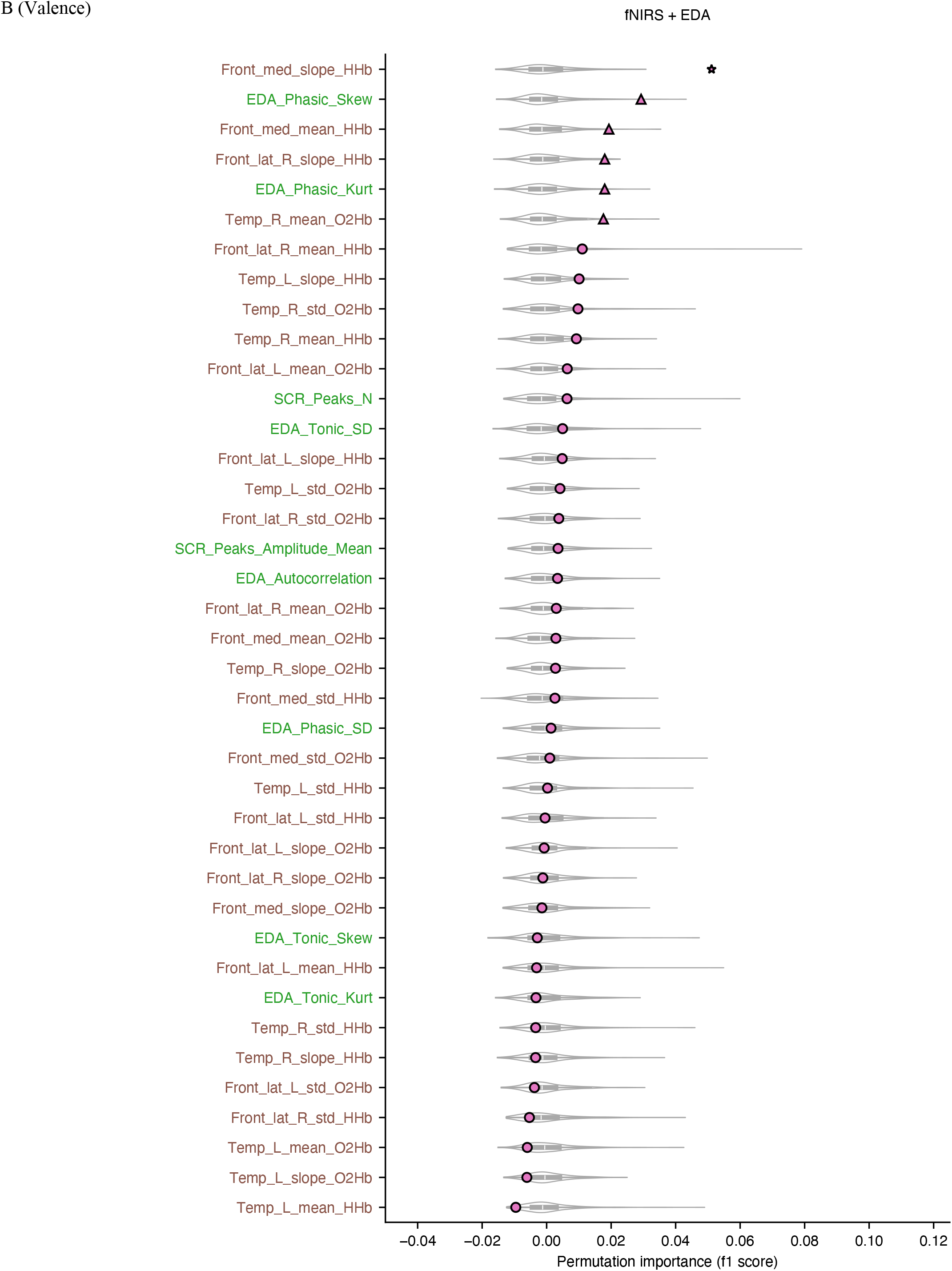
Permutation importance for emotion classification using the fNIRS + EDA feature set. Permutation importance was computed with macro-averaged F1 as the performance metric. A higher permutation importance score indicates a greater contribution to emotion classification. Violin plots show the null distributions generated by the permutation test. Pink dots indicate the median of the 20 permutation-importance estimates obtained from repeated grouped 5-fold cross-validation. The marker shape denotes the significance of the observed permutation importance relative to the null distribution (circle: *p* ≥ 0.05; triangle: *p* < 0.05; star: *p* < 0.005). The color of the y-axis labels indicates the modality to which each feature belongs (brown: fNIRS; green: EDA). (A) Arousal. (B) Valence. front: prefrontal, lat: lateral, med: medial, temp: temporal, HHb: deoxygenated hemoglobin signal, O2Hb: oxygenated hemoglobin signal, L: left, R: right, Peaks_N: the number of peaks, skew: skewness, kurt: kurtosis, SD: standard deviation (std), SCR: skin conductance response (phasic signal).

Fig. 5 indicates that only a subset of features substantially contributed to classification in both the arousal and valence tasks. Moreover, EDA-derived features showed more pronounced importance for arousal, whereas fNIRS-derived features tended to exhibit relatively higher importance for valence.

Permutation-importance results for the other feature sets are provided in the Supplementary Material (Figs. S4A–L).

## IV. Discussion

In this study, we investigated whether augmenting fNIRS— used to capture central nervous system activity—with two peripheral physiological modalities, EDA and PPG, improves high/low classification of the fundamental affective dimensions of arousal and valence. The highest performance for both arousal and valence was obtained when fNIRS was combined with EDA. The results offer empirical evidence supporting the effectiveness of combining fNIRS with peripheral physiological data for classifying emotions.

An important aspect of the present work is that the benefit of multimodal sensing was demonstrated without relying on complex modeling pipelines. Specifically, we intentionally restricted the input representation to commonly used, easily computed features and employed a classical classifier, without feature selection, extensive feature engineering, or hyperparameter optimization. This design choice prioritizes reproducibility and transparency of the analysis pipeline.

Building on these results, future work may explore whether more advanced approaches—such as deep learning or sensor-level fusion methods—can further improve performance, while carefully balancing accuracy gains against added complexity and reduced interpretability.

Across both affective dimensions, combining fNIRS with peripheral signals improved performance relative to fNIRS alone, indicating that multimodal integration is beneficial. The valence results are particularly informative. Although EDA and PPG did not yield statistically significant performance when used alone for valence classification, combining each of them with fNIRS improved performance beyond fNIRS alone. This pattern suggests that peripheral features may provide complementary information that is not sufficient by itself to support reliable valence discrimination, but becomes informative when integrated with central signals. This interpretation is consistent with the permutation-importance analysis: for valence classification, top-ranked features included contributions from both fNIRS and peripheral modalities (fNIRS + EDA in Fig. 5B; fNIRS + PPG in Fig. S4J).

In contrast, the feature-importance patterns differed for arousal. When fNIRS was combined with EDA, EDA features dominated the most informative ranks, whereas when fNIRS was combined with PPG, fNIRS features were more prominent. Taken together, these results suggest that arousal can often be inferred from a single modality, whereas valence may require complementary information from both central and peripheral measurements.

The combination of EDA and PPG provided limited benefit. For arousal, combining EDA and PPG yielded lower performance than EDA alone, and for valence the combination did not achieve statistically significant performance (consistent with the unimodal results). This indicates redundancy between these two peripheral modalities in the present setting, and suggests that combining EDA and PPG without central measurements may be insufficient for robust emotion classification. Related work has similarly reported limited gains from combining these modalities [33].

Unlike the peripheral modalities, fNIRS alone achieved statistically significant classification for both arousal and valence, indicating that central hemodynamic signals contain information relevant to discriminating these affective dimensions. This result is consistent with prior work demonstrating subject-independent arousal/valence classification using fNIRS [20]. More broadly, these findings support the view that multimodal affective computing frameworks centered on central nervous system measurements can be effective, and highlight the practical value of fNIRS for real-world settings due to its relative robustness to electromagnetic noise and certain motion-related artifacts.

EDA showed strong performance primarily for arousal, consistent with the common characterization of EDA as a reliable index of arousal but less informative for valence [34]. A recent survey has also quantitatively reported that arousal tends to be easier to decode than valence from EDA signals [35], in line with our observations. At the same time, some studies have reported subject-independent classification of both arousal and valence using EDA alone [36], [37].

Resolving these discrepancies likely requires systematic investigation of factors such as experimental paradigms, labeling protocols, feature definitions, and subject populations. At minimum, the present results reinforce the utility of EDA for arousal-related classification.

PPG achieved statistically significant performance for arousal, but with lower accuracy than the other modalities and combinations considered. Prior work has reported high classification performance for both arousal and valence using PPG [38], which is not fully consistent with our findings. Several factors may contribute to this discrepancy (e.g., differences in stimuli, recording conditions, and preprocessing). Importantly, our pipeline intentionally used a simple, toolbox-based feature set without feature selection to maximize reproducibility and transparency. It is plausible that more carefully tailored PPG features and quality-control procedures could improve performance; however, optimizing PPG-only classification was not a primary objective of this study.

This study has three limitations. First, the analysis pipeline was intentionally constrained to simple features and a classical classifier to prioritize reproducibility and transparency. While this design supports transparent benchmarking, future work should evaluate whether modern sensor-level fusion methods [12] and deep learning can provide meaningful improvements. Second, the dataset comprised predominantly young university students with a male-skewed gender distribution (male:female = 2:1). Therefore, it remains to be tested whether the observed performance patterns generalize to cohorts with balanced gender ratios or different age ranges. Previous research on affect recognition utilizing EDA, electromyography (EMG), and heart rate indicates superior performance in young females compared to young males, as well as enhanced performance in older individuals relative to younger cohorts [39]. These findings raise the possibility that performance could differ—potentially improve—in more diverse populations, which warrants future validation. Third, peripheral physiology was represented only by EDA and PPG in the present study. Other peripheral modalities—such as electrocardiography, skin temperature, electrooculography, and EMG—may provide additional complementary information. Future work should systematically evaluate which peripheral signals are most synergistic with fNIRS for subject-independent emotion classification.

## V. Conclusion

This study investigated whether subject-independent emotion classification can benefit from multimodal data that combine central nervous system activity measured by fNIRS with peripheral physiological signals, and in particular whether such multimodal sensing improves performance relative to fNIRS alone. We observed higher classification performance when fNIRS was combined with either EDA or PPG, with the strongest performance obtained when fNIRS was combined with EDA. These results underscore the importance of integrating fNIRS with peripheral physiological measurements for emotion classification. Given the relative robustness of fNIRS to environmental electromagnetic noise and certain motion-related artifacts, fNIRS-centered multimodal sensing may be a promising direction for enabling affective computing in more naturalistic environments.

## Acknowledgment

S. I. would like to thank M. K. for assistance with music-video selection. The authors used ChatGPT 5.4 (Thinking) developed by OpenAI and Paperpal developed by Editage for manuscript preparation support, including translation from Japanese to English, paraphrasing, proofreading, and partial image generation for Fig. 2.

