## Supplementary Materials for "Improving Emotion Classification by Combining fNIRS-Derived Hemodynamic Responses with Peripheral Physiological Signals"

Table S1 Forty Japanese popular music videos used as candidate stimuli. Each selected video ID corresponds to the numbering shown in Figs. 1 and 3. The start time of each extracted clip is listed under Highlight Start (m:ss). When the artist name or original song title was written in Japanese script, it was romanized and presented as romanized form (original Japanese script).

| Selected Video ID | Emotion Label | Artist | Title | YouTube Link | Highlight Start (m:ss) |
|---|---|---|---|---|---|
| 2 | high arousal | SANDAIME J SOUL BROTHERS from EXILE TRIBE (三代目 J SOUL BROTHERS from EXILE TRIBE) | R.Y.U.S.E.I. | https://www.youtube.com/watch?v=4-Gw0TAM6-Q | 1:00 |
| | | AKB48 | Flying Get (フライングゲット) | https://www.youtube.com/watch?v=WdhMjzfg6-k&list=RDWdhMjzfg6-k&start_radio=1 | 2:50 |
| | | YOASOBI | Idol (アイドル) | https://www.youtube.com/watch?v=ZRtdQ81jPUQ | 0:18 |
| | | Ayaka (絢香) | Nijiiro (にじいろ) | https://www.youtube.com/watch?v=ia0lAgfhbAo | 0:17 |
| | | Golden Bomber (ゴールデンボンバー) | Memeshikute (女々しくて) | https://www.youtube.com/watch?v=BC9P3DSZu0A | 3:20 |
| | | Mary's Blood | Pegasus Fantasy (ペガサス幻想) | https://www.youtube.com/watch?v=M-XG-RW79eE | 2:45 |
| | | Ling tosite sigure (凛として時雨) | unravel | https://www.youtube.com/watch?v=Fve_lHIPa-I | 2:27 |
| 3 | high arousal | T.M.Revolution | HOT LIMIT | https://www.youtube.com/watch?v=vBmU5v2EyxM | 2:35 |
| | | GReeeeN | Aiueongaku (あいうえおんがく) | https://www.youtube.com/watch?v=ADqLniFy6ww | 0:33 |
| | | ORANGE RANGE | SUSHI Tabetai (SUSHI 食べたい) | https://www.youtube.com/watch?v=u7adbYXou8Q | 0:00 |
| | | Ado | Usseewa (うっせぇわ) | https://www.youtube.com/watch?v=Qp3b-RXtz4w | 1:13 |
| | | Keyakizaka46 (欅坂46) | Fukyouwaon (不協和音) | https://www.youtube.com/watch?v=gfzuzDrVRVM | 3:15 |
| 1 | high arousal | Keyakizaka46 (欅坂46) | Garasu Wo Ware! (ガラスを割れ！) | https://www.youtube.com/watch?v=A2k6ZO6B0A8 | 0:33 |
| 12 | low valence | ALI PROJECT | Kyomu Densen (凶夢伝染) | https://www.youtube.com/watch?v=We_uZ7-_HmM | 2:59 |
| | | natori (なとり) | Shokutaku (食卓) | https://www.youtube.com/watch?v=Ph5xcTn_I1I | 2:04 |
| | | CHANMINA (ちゃんみな) | Bijin (美人) | https://www.youtube.com/watch?v=r2bHEcxB5kY | 0:10 |
| | | Shimamiya Eiko (島みやえい子) | Higurashi no Naku Koro ni (ひぐらしのなく頃に) | https://www.youtube.com/watch?v=V9nV4SHbcBA | 0:26 |
| | | EGOIST | Namae No Nai Kaibutsu (名前のない怪物) | https://www.youtube.com/watch?v=qiX5DI--8bg | 4:19 |
| | | Shinsei Kamattechan | Boku no Sensou | https://www.youtube.com/watch?v=goXKlOozyx8 | 3:17 |

| | | | | | |
|---|---|---|---|---|---|
| | | (神聖かまってちゃん) | (僕の戦争) | | |
| 10 | low valence | amazarashi | Seasons die one after another (季節は次々死んでいく) | https://www.youtube.com/watch?v=wtJcLWeY114 | 4:29 |
| 8 | high valence | Spitz (スピッツ) | Cherry (チェリー) | https://www.youtube.com/watch?v=Eze6-eHmtJg | 0:21 |
| | | Yonezu Kenshi (米津玄師) | Canary (カナリヤ) | https://www.youtube.com/watch?v=JAMNqRBL_CY | 0:13 |
| 6 | low arousal | Kamishiraishi Moka, HY (上白石萌歌、HY) | 366 Days (366日) | https://www.youtube.com/watch?v=HIB8RBhPkBA | 0:32 |
| | | KOBUKURO (コブクロ) | Tsubomi (蕾) | https://www.youtube.com/watch?v=WPH1BLHKOJE | 0:15 |
| | | Saito Kazuyoshi (斉藤和義) | Uta Utai no Ballad (歌うたいのバラッド) | https://www.youtube.com/watch?v=9G82-cb5zUk | 0:30 |
| | | Fujiwara Sakura (藤原さくら) | Soup | https://www.youtube.com/watch?v=cd-qx2lF9Hg | 0:00 |
| 7 | high valence | Suda Masaki (菅田将暉) | Niji (虹) | https://www.youtube.com/watch?v=hkBbUf4oGfA | 1:28 |
| | | Ohara Sakurako (大原櫻子) | Hitomi (瞳) | https://www.youtube.com/watch?v=TojGErB7pHY | 0:04 |
| 9 | high valence | SEKAI NO OWARI | Sasanqua (サザンカ) | https://www.youtube.com/watch?v=249YdrcCL0Y | 0:28 |
| | | Matsutoya Yumi (松任谷由実) | Haru Yo, Koi (春よ、来い) | https://www.youtube.com/watch?v=qX7pFYH9O04 | 1:27 |
| 4 | low arousal | AI | Okuribito (おくりびと) | https://www.youtube.com/watch?v=XHenwMKcSL0 | 0:05 |
| | | LAMP IN TERREN | Calm Down (カームダウン) | https://www.youtube.com/watch?v=0y8k3ySffOA | 0:00 |
| 5 | low arousal | Ms.OOJA | HIKARI | https://www.youtube.com/watch?v=BLEnWCSNG8Q | 0:20 |
| 11 | low valence | Kikuo | Aishite Aishite Aishite (愛して愛して愛して) | https://www.youtube.com/watch?v=NTrm_idbhUk | 1:53 |
| | | Nakashima Mika (中島美嘉) | Boku Ga Shinou To Omottanowa (僕が死のうと思ったのは) | https://www.youtube.com/watch?v=ZH-amYi7_o4 | 0:06 |
| | | Daoko | Rinko (燐光) | https://www.youtube.com/watch?v=rLYAStnZDtY | 0:59 |
| | | ReoNa | Ikiterudakede Eraiyo (生きてるだけでえらいよ) | https://www.youtube.com/watch?v=6YIf6UwoYrM | 0:07 |
| | | Onitsuka Chihiro (鬼束ちひろ) | Gekkou (月光) | https://www.youtube.com/watch?v=iyw6-KVmgow | 2:10 |
| | | Hirai Ken (平井堅) | Kokuhaku (告白) | https://www.youtube.com/watch?v=_o10f0CMTvU | 0:00 |

| | | | | |
|---|---|---|---|---|
| | Yuuri (優里) | Merry-go-round (メリーゴーランド) | https://www.youtube.com/watch?v=eWeSqrRk-gs | 2:12 |

Table S2 Twenty-seven features computed from the PPG signal.

| Feature Domain | Feature Abbreviations (NeuroKit2) | Description of Each Feature (Excerpted from NeuroKit2) |
|---|---|---|
| time-domain | PPG_Rate_Mean | PPG_Rate_Mean: The mean PPG rate. |
| time-domain | HRV_MeanNN | MeanNN: The mean of the RR intervals. |
| time-domain | HRV_SDNN | SDNN: The standard deviation of the RR intervals. |
| time-domain | HRV_RMSSD | RMSSD: The square root of the mean of the squared successive differences between adjacent RR intervals. It is equivalent (although on another scale) to SD1, and therefore it is redundant to report correlations with both (Ciccone, 2017). |
| time-domain | HRV_SDSD | SDSD: The standard deviation of the successive differences between RR intervals. |
| time-domain | HRV_CVNN | CVNN: The standard deviation of the RR intervals (SDNN) divided by the mean of the RR intervals (MeanNN). |
| time-domain | HRV_CVSD | CVSD: The root mean square of successive differences (RMSSD) divided by the mean of the RR intervals (MeanNN). |
| time-domain | HRV_MedianNN | MedianNN: The median of the RR intervals. |
| time-domain | HRV_MadNN | MadNN: The median absolute deviation of the RR intervals. |
| time-domain | HRV_MCVNN | MCVNN: The median absolute deviation of the RR intervals (MadNN) divided by the median of the RR intervals (MedianNN). |
| time-domain | HRV_IQRNN | IQRNN: The interquartile range (IQR) of the RR intervals. |
| time-domain | HRV_SDRMSSD | SDRMSSD: SDNN / RMSSD, a time-domain equivalent for the low Frequency-to-High Frequency (LF/HF) Ratio (Sollers et al., 2007). |
| time-domain | HRV_Prc20NN | Prc20NN: The 20th percentile of the RR intervals (Han, 2017; Hovsepian, 2015). |
| time-domain | HRV_Prc80NN | Prc80NN: The 80th percentile of the RR intervals (Han, 2017; Hovsepian, 2015). |
| time-domain | HRV_pNN20 | pNN20: The percentage of absolute differences in successive RR intervals greater than 20 ms (Mietus et al., 2002). |
| time-domain | HRV_MinNN | MinNN: The minimum of the RR intervals (Parent, 2019; Subramaniam, 2022). |
| time-domain | HRV_MaxNN | MaxNN: The maximum of the RR intervals (Parent, 2019; Subramaniam, 2022). |
| time-domain | HRV_HTI | HTI: The HRV triangular index, measuring the total number of RR intervals divided by the height of the RR intervals histogram. |
| time-domain | HRV_TINN | TINN: A geometrical parameter of the HRV, or more specifically, the baseline width of the RR intervals distribution obtained by triangular interpolation, where the error of least squares determines the triangle. It is an approximation of the RR interval distribution. |
| frequency-domain | HRV_LF | HRV_LF: The spectral power of low frequencies (by default, .04 to .15 Hz). |
| frequency-domain | HRV_HF | HRV_HF: The spectral power of high frequencies (by default, .15 to .4 Hz). |
| frequency-domain | HRV_VHF | HRV_VHF: The spectral power of very high frequencies (by default, .4 to .5 Hz). |
| frequency-domain | HRV_TP | HRV_TP: The total spectral power. |
| frequency-domain | HRV_LFHF | HRV_LFHF: The ratio obtained by dividing the low frequency power by the high frequency power. |

| | | |
|---|---|---|
| frequency-domain | HRV_LFn | HRV_LFn: The normalized low frequency, obtained by dividing the low frequency power by the total power. |
| frequency-domain | HRV_HFn | HRV_HFn: The normalized high frequency, obtained by dividing the low frequency power by the total power. |
| frequency-domain | HRV_LnHF | HRV_LnHF: The log transformed HF. |

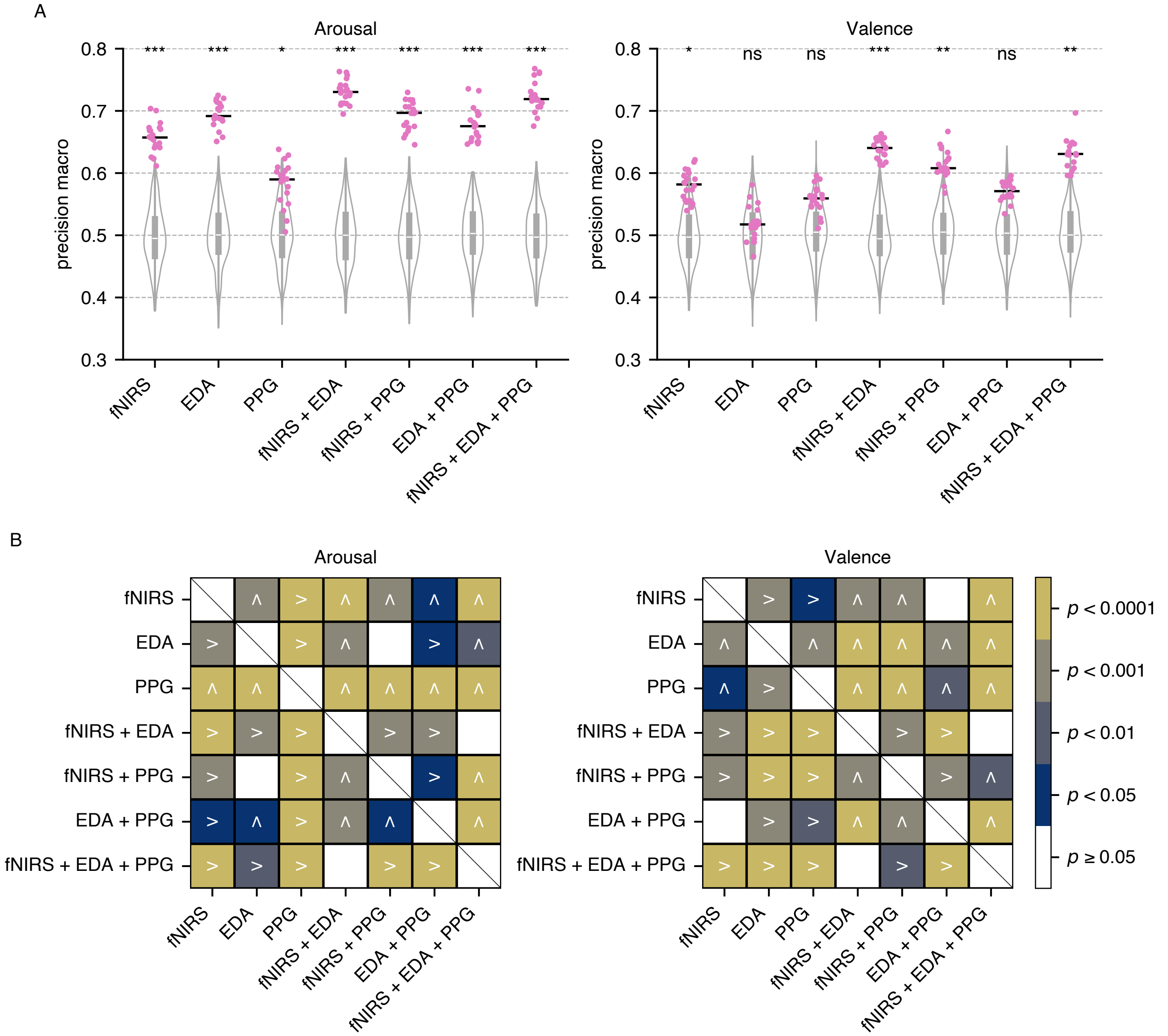


Fig. S1. Results of emotion classification. (A) Macro-averaged precision scores for each feature set. Pink dots denote the 20 performance estimates obtained from repeated grouped 5-fold cross-validation, and the horizontal black line indicates the median across repetitions. Violin plots show the null distributions generated by the permutation test. Asterisks indicate significance levels of the observed performance relative to the null distribution (ns: $p \geq 0.05$; * $p < 0.05$; ** $p < 0.01$; *** $p < 0.005$). (B) Pairwise comparisons of macro-averaged precision scores between feature sets; colors indicate significance levels from the two-sided Wilcoxon signed-rank test. The symbol “>” denotes that the feature set in the row outperformed that in the column, whereas the “∧” symbol denotes the opposite (i.e., the feature set in the column outperformed that in the row).

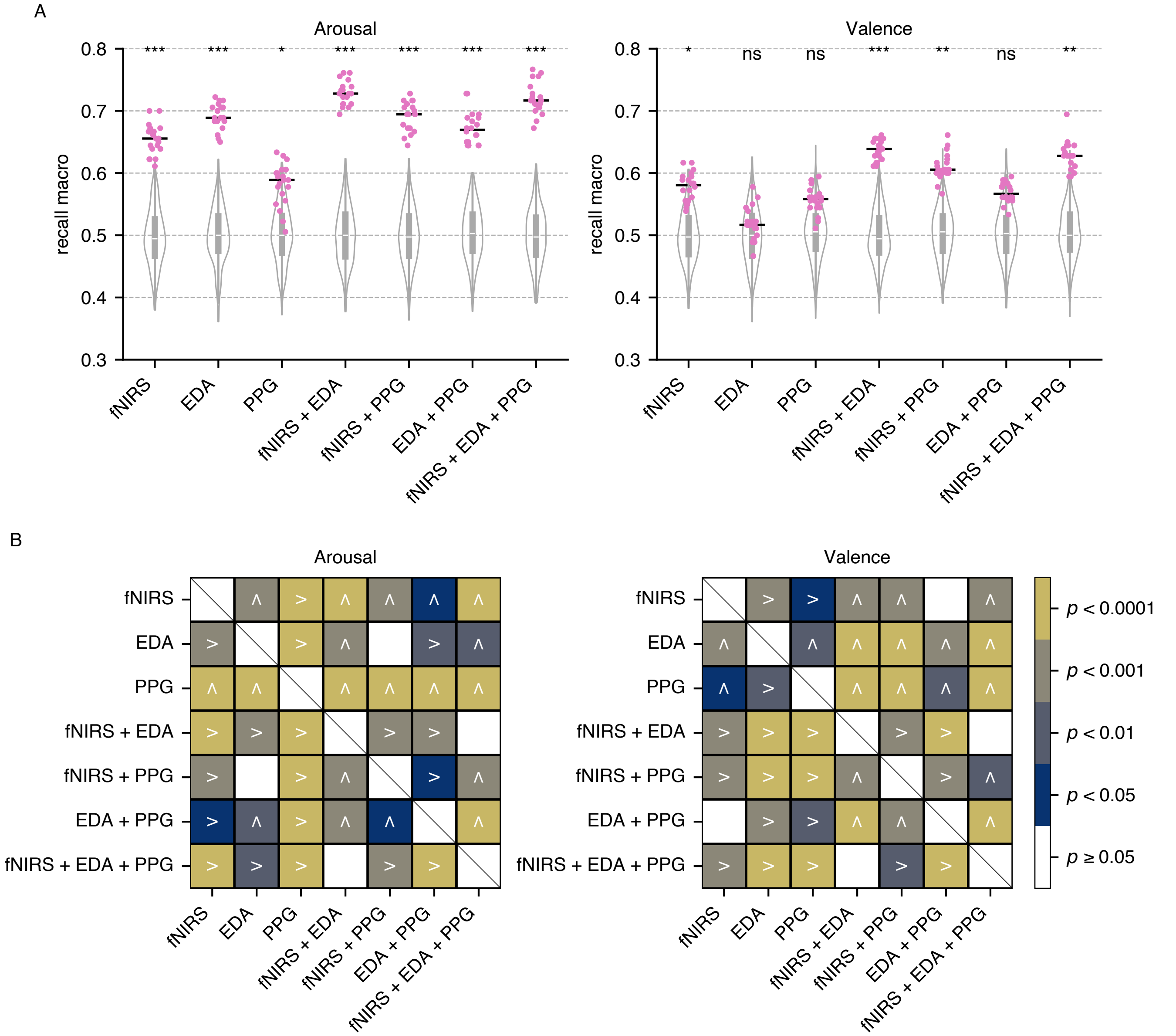


Fig. S2. Results of emotion classification. (A) Macro-averaged recall scores for each feature set. Pink dots denote the 20 performance estimates obtained from repeated grouped 5-fold cross-validation, and the horizontal black line indicates the median across repetitions. Violin plots show the null distributions generated by the permutation test. Asterisks indicate significance levels of the observed performance relative to the null distribution (ns: $p \geq 0.05$; * $p < 0.05$; ** $p < 0.01$; *** $p < 0.005$). (B) Pairwise comparisons of macro-averaged recall scores between feature sets; colors indicate significance levels from the two-sided Wilcoxon signed-rank test. The symbol “>” denotes that the feature set in the row outperformed that in the column, whereas the “∧” symbol denotes the opposite (i.e., the feature set in the column outperformed that in the row).

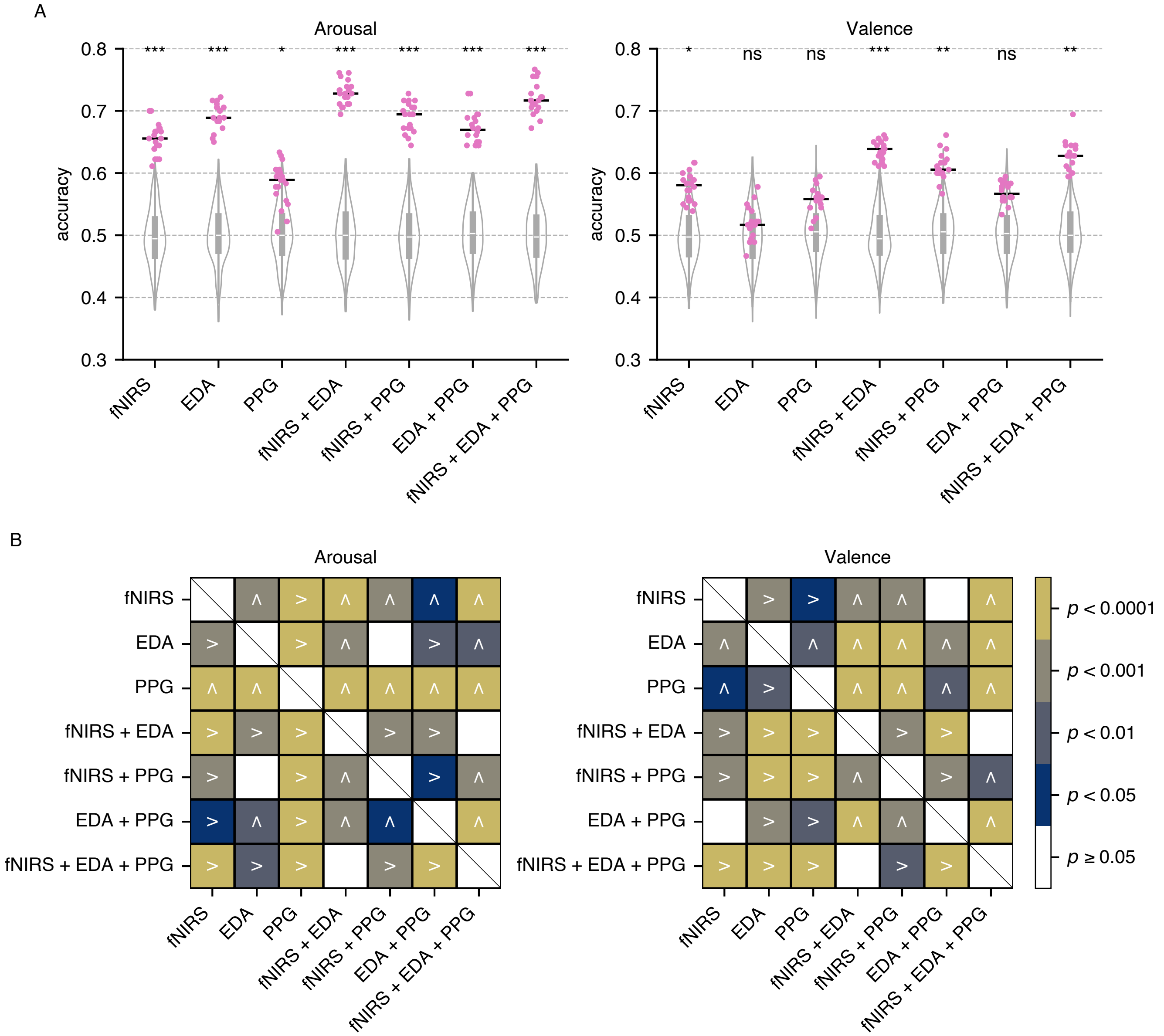


Fig. S3. Results of emotion classification. (A) Accuracy scores for each feature set. Pink dots denote the 20 performance estimates obtained from repeated grouped 5-fold cross-validation, and the horizontal black line indicates the median across repetitions. Violin plots show the null distributions generated by the permutation test. Asterisks indicate significance levels of the observed performance relative to the null distribution (ns: $p \geq 0.05$; * $p < 0.05$; ** $p < 0.01$; *** $p < 0.005$). (B) Pairwise comparisons of accuracy scores between feature sets; colors indicate significance levels from the two-sided Wilcoxon signed-rank test. The symbol ">" denotes that the feature set in the row outperformed that in the column, whereas the "∧" symbol denotes the opposite (i.e., the feature set in the column outperformed that in the row).

A (Arousal)

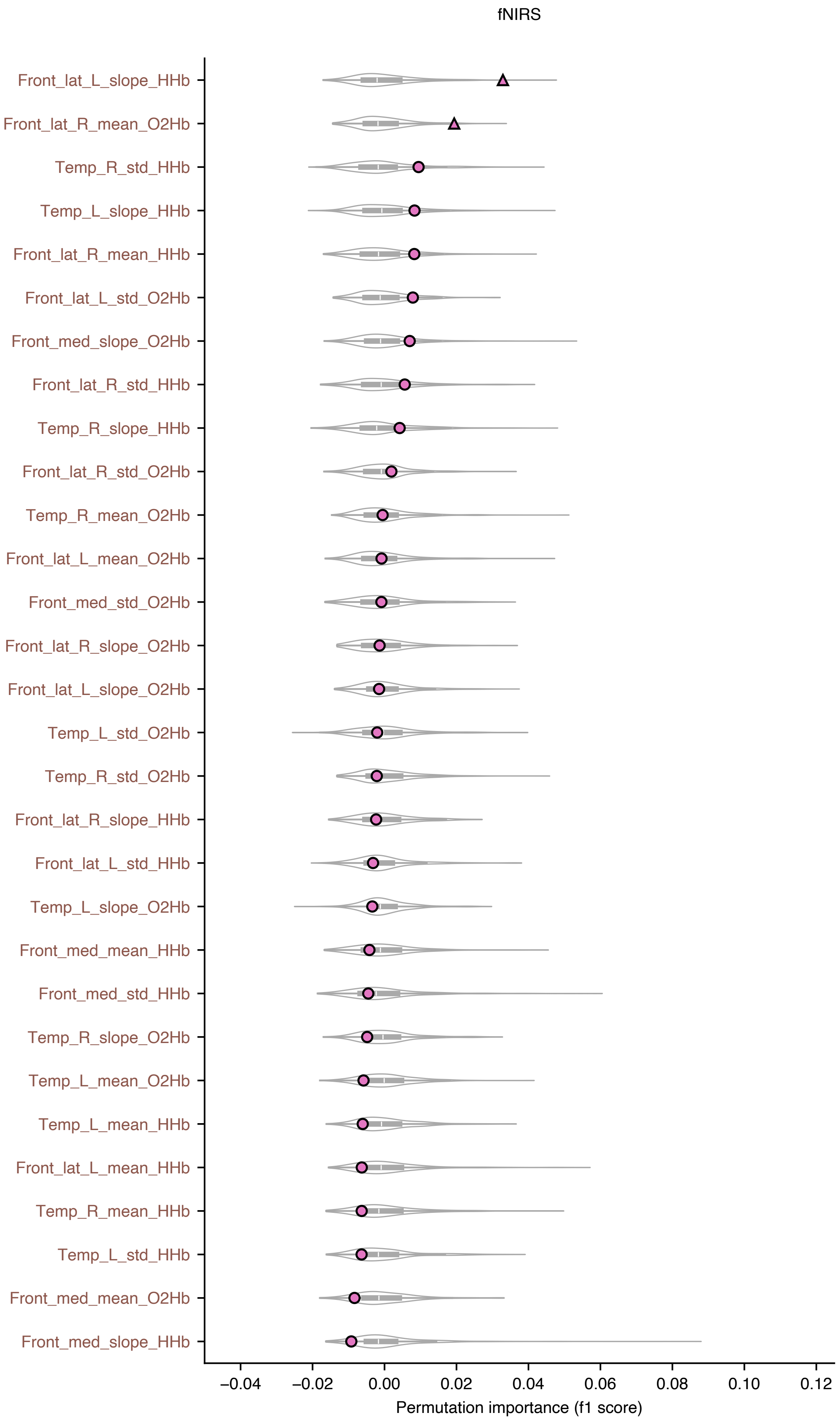


Fig. S4. Permutation importance for emotion classification using individual feature sets. Permutation importance was computed with macro-averaged F1 as the performance metric. A higher permutation importance score indicates a greater contribution to emotion classification. Violin plots show the null distributions generated by the permutation test. Pink dots indicate the median of the 20 permutation-importance estimates obtained from repeated grouped 5-fold cross-validation. The marker shape denotes the significance of the observed permutation importance relative to the null distribution (circle: $p \geq 0.05$; triangle: $p < 0.05$; square: $p < 0.01$; star: $p < 0.005$). The color of the y-axis labels indicates the modality to which each feature belongs (brown: fNIRS; green: EDA; cyan: PPG). front: prefrontal, lat: lateral, med: medial, temp: temporal, HHb: deoxygenated hemoglobin signal, O2Hb: oxygenated hemoglobin signal, L: left, R: right, Peaks_N: the number of peaks, skew: skewness, kurt: kurtosis, SD: standard deviation (std), SCR: skin conductance response (phasic signal).

(A) fNIRS (Arousal).
(B) EDA (Arousal).
(C) PPG (Arousal).
(D) fNIRS + PPG (Arousal).
(E) EDA + PPG (Arousal).
(F) fNIRS + EDA + PPG (Arousal).
(G) fNIRS (Valence).
(H) EDA (Valence).
(I) PPG (Valence).
(J) fNIRS + PPG (Valence).
(K) EDA + PPG (Valence).
(L) fNIRS + EDA + PPG (Valence).

B (Arousal)

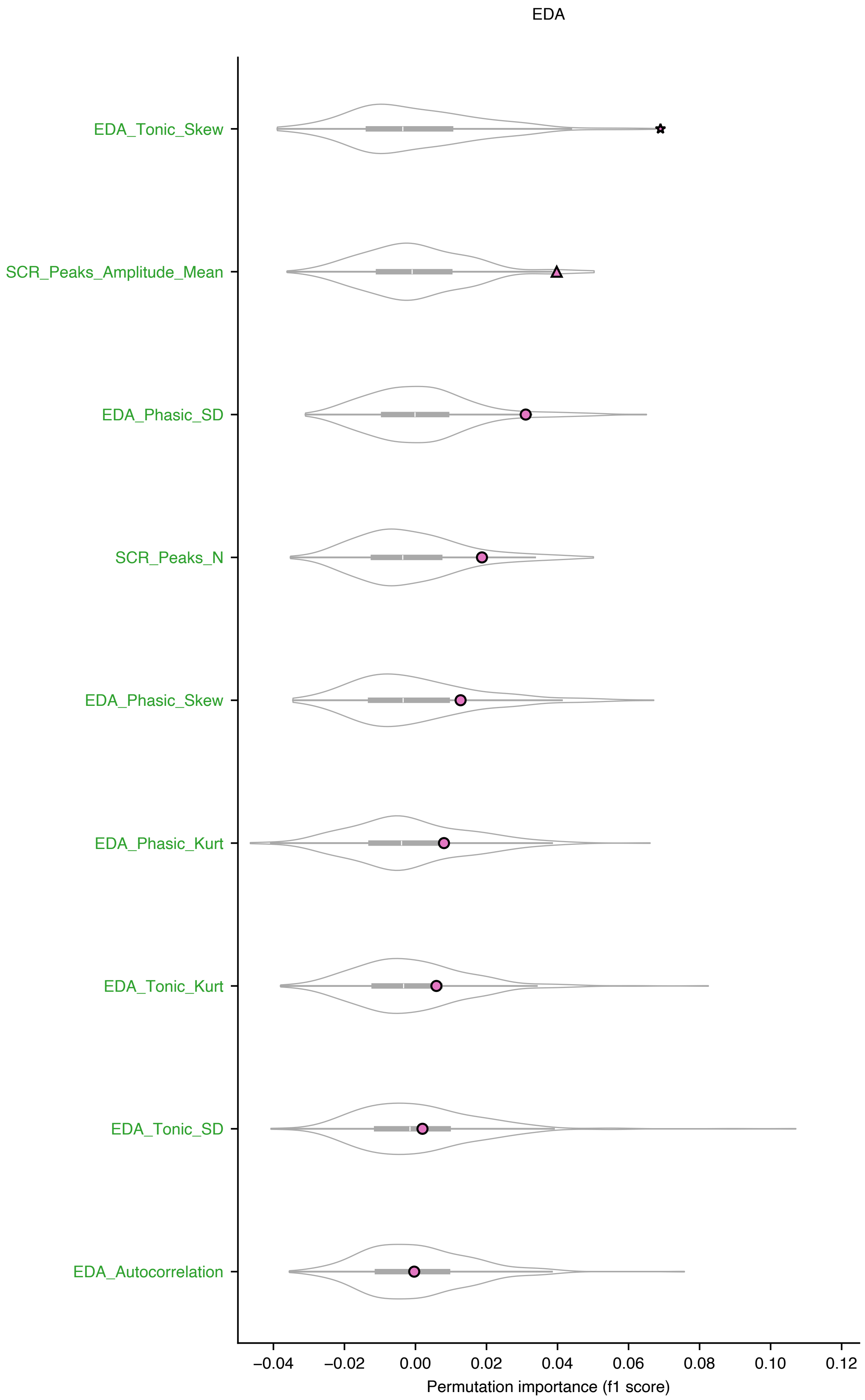

EDA
EDA_Tonic_Skew
SCR_Peaks_Amplitude_Mean
EDA_Phasic_SD
SCR_Peaks_N
EDA_Phasic_Skew
EDA_Phasic_Kurt
EDA_Tonic_Kurt
EDA_Tonic_SD
EDA_Autocorrelation
−0.04
−0.02
0.00
0.02
0.04
0.06
0.08
0.10
0.12
Permutation importance (f1 score)

C (Arousal)

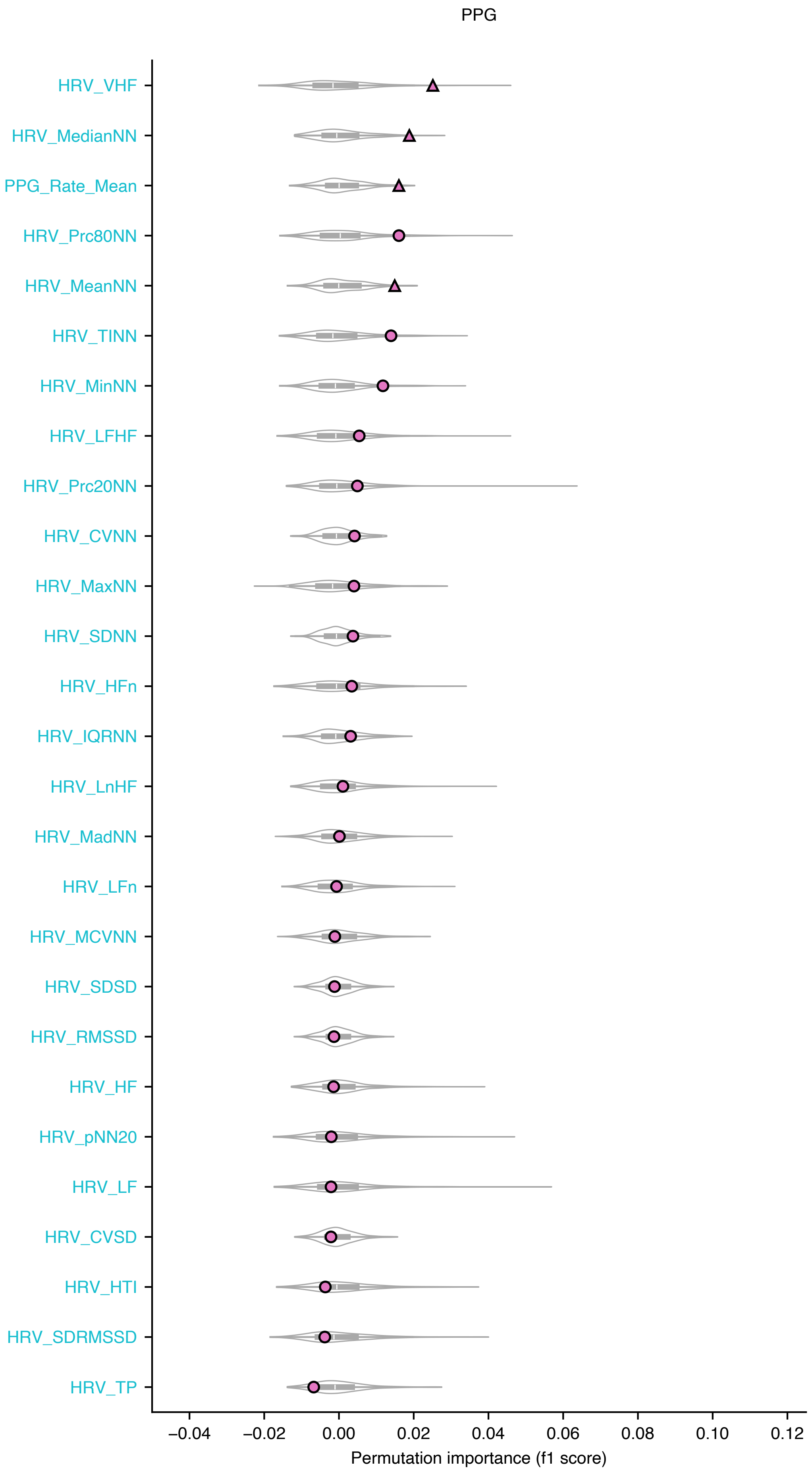
PPG
HRV_VHF
HRV_MedianNN
PPG_Rate_Mean
HRV_Prc80NN
HRV_MeanNN
HRV_TINN
HRV_MinNN
HRV_LFHF
HRV_Prc20NN
HRV_CVNN
HRV_MaxNN
HRV_SDNN
HRV_HFn
HRV_IQRNN
HRV_LnHF
HRV_MadNN
HRV_LFn
HRV_MCVNN
HRV_SDSD
HRV_RMSSD
HRV_HF
HRV_pNN20
HRV_LF
HRV_CVSD
HRV_HTI
HRV_SDRMSSD
HRV_TP
−0.04
−0.02
0.00
0.02
0.04
0.06
0.08
0.10
0.12
Permutation importance (f1 score)

D (Arousal)

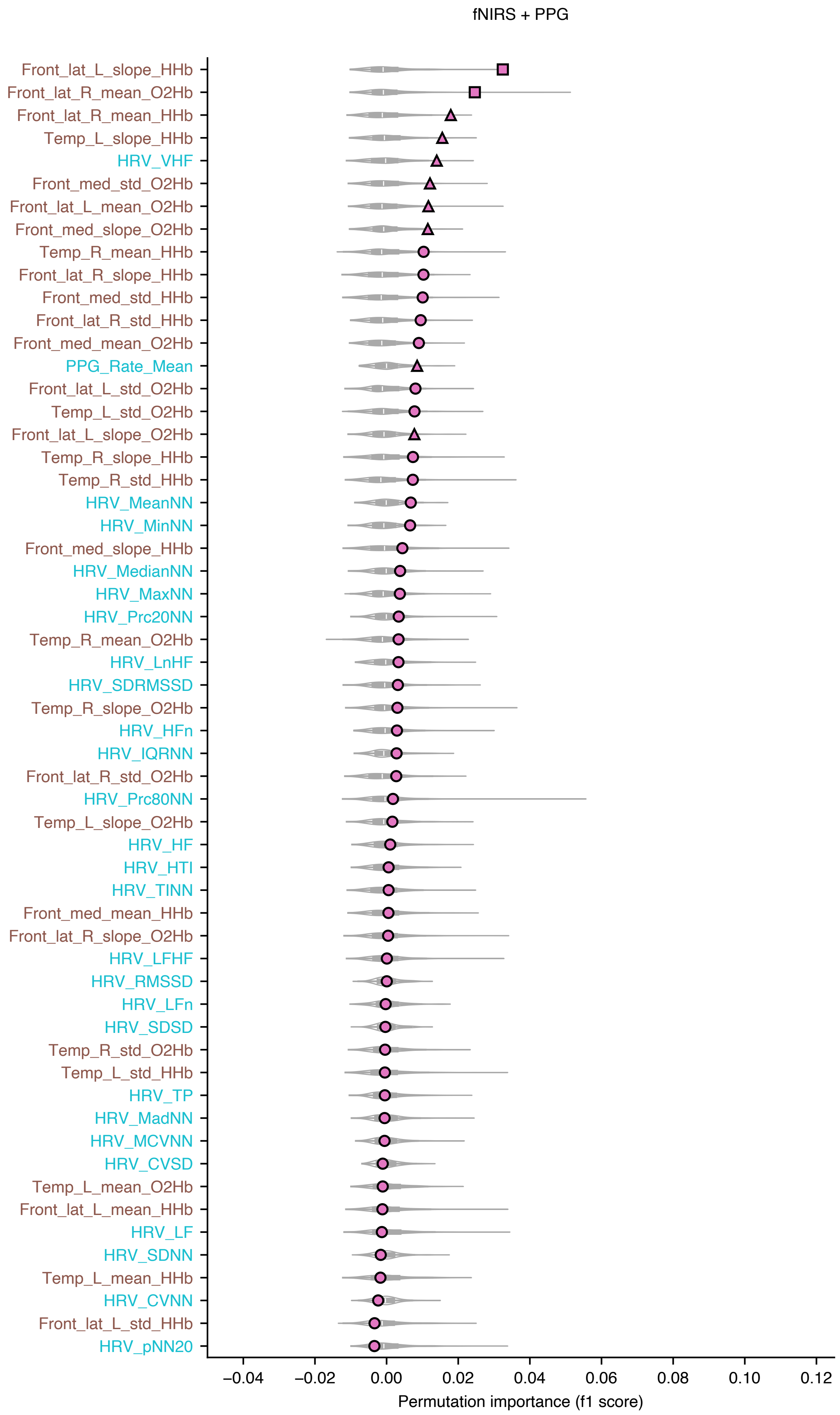

fNIRS + PPG
Front_lat_L_slope_HHb
Front_lat_R_mean_O2Hb
Front_lat_R_mean_HHb
Temp_L_slope_HHb
HRV_VHF
Front_med_std_O2Hb
Front_lat_L_mean_O2Hb
Front_med_slope_O2Hb
Temp_R_mean_HHb
Front_lat_R_slope_HHb
Front_med_std_HHb
Front_lat_R_std_HHb
Front_med_mean_O2Hb
PPG_Rate_Mean
Front_lat_L_std_O2Hb
Temp_L_std_O2Hb
Front_lat_L_slope_O2Hb
Temp_R_slope_HHb
Temp_R_std_HHb
HRV_MeanNN
HRV_MinNN
Front_med_slope_HHb
HRV_MedianNN
HRV_MaxNN
HRV_Prc20NN
Temp_R_mean_O2Hb
HRV_LnHF
HRV_SDRMSSD
Temp_R_slope_O2Hb
HRV_HFn
HRV_IQRNN
Front_lat_R_std_O2Hb
HRV_Prc80NN
Temp_L_slope_O2Hb
HRV_HF
HRV_HTI
HRV_TINN
Front_med_mean_HHb
Front_lat_R_slope_O2Hb
HRV_LFHF
HRV_RMSSD
HRV_LFn
HRV_SDSD
Temp_R_std_O2Hb
Temp_L_std_HHb
HRV_TP
HRV_MadNN
HRV_MCVNN
HRV_CVSD
Temp_L_mean_O2Hb
Front_lat_L_mean_HHb
HRV_LF
HRV_SDNN
Temp_L_mean_HHb
HRV_CVNN
Front_lat_L_std_HHb
HRV_pNN20
−0.04
−0.02
0.00
0.02
0.04
0.06
0.08
0.10
0.12
Permutation importance (f1 score)

E (Arousal)

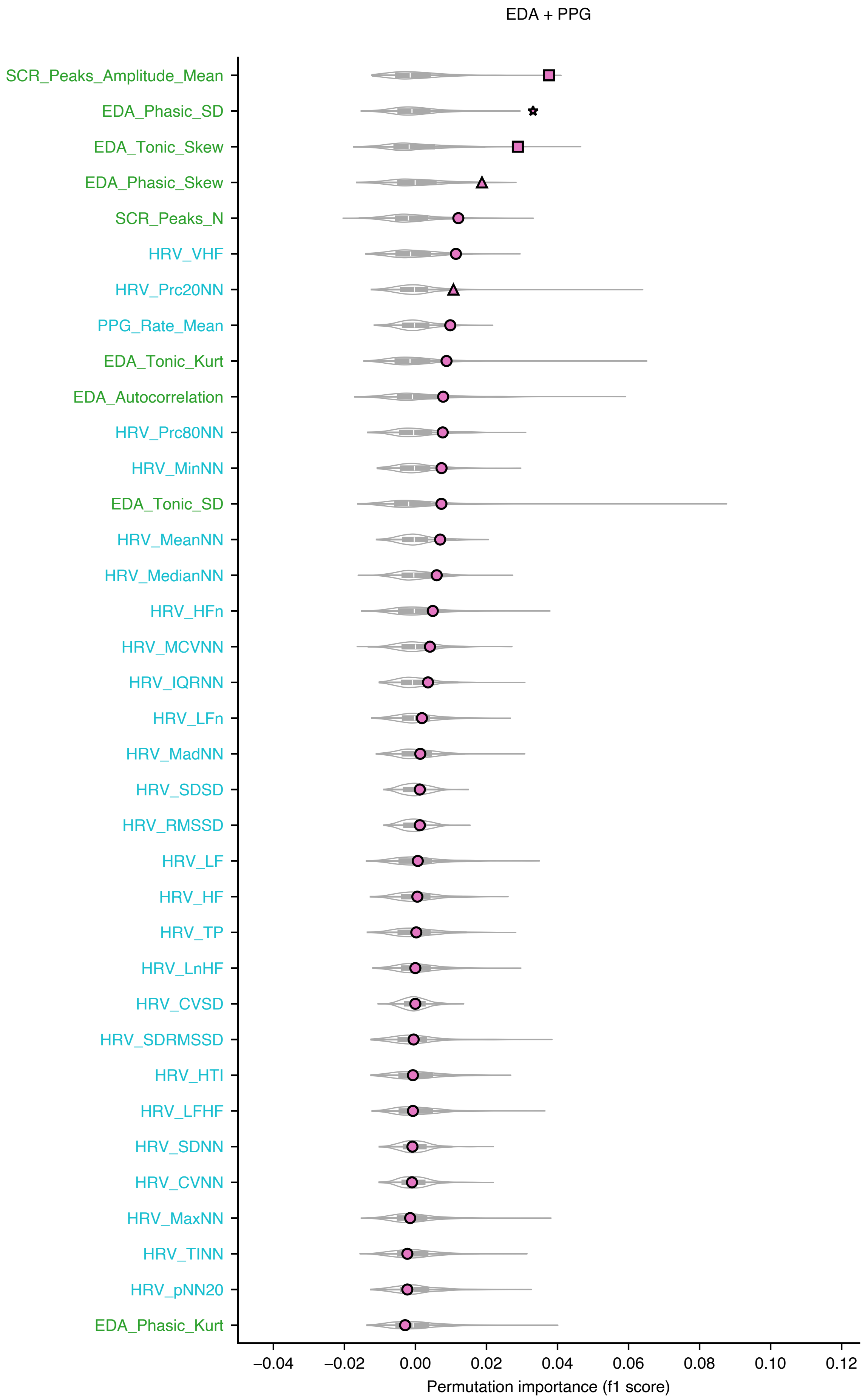
EDA + PPG
SCR_Peaks_Amplitude_Mean
EDA_Phasic_SD
EDA_Tonic_Skew
EDA_Phasic_Skew
SCR_Peaks_N
HRV_VHF
HRV_Prc20NN
PPG_Rate_Mean
EDA_Tonic_Kurt
EDA_Autocorrelation
HRV_Prc80NN
HRV_MinNN
EDA_Tonic_SD
HRV_MeanNN
HRV_MedianNN
HRV_HFn
HRV_MCVNN
HRV_IQRNN
HRV_LFn
HRV_MadNN
HRV_SDSD
HRV_RMSSD
HRV_LF
HRV_HF
HRV_TP
HRV_LnHF
HRV_CVSD
HRV_SDRMSSD
HRV_HTI
HRV_LFHF
HRV_SDNN
HRV_CVNN
HRV_MaxNN
HRV_TINN
HRV_pNN20
EDA_Phasic_Kurt
−0.04
−0.02
0.00
0.02
0.04
0.06
0.08
0.10
0.12
Permutation importance (f1 score)

F (Arousal)

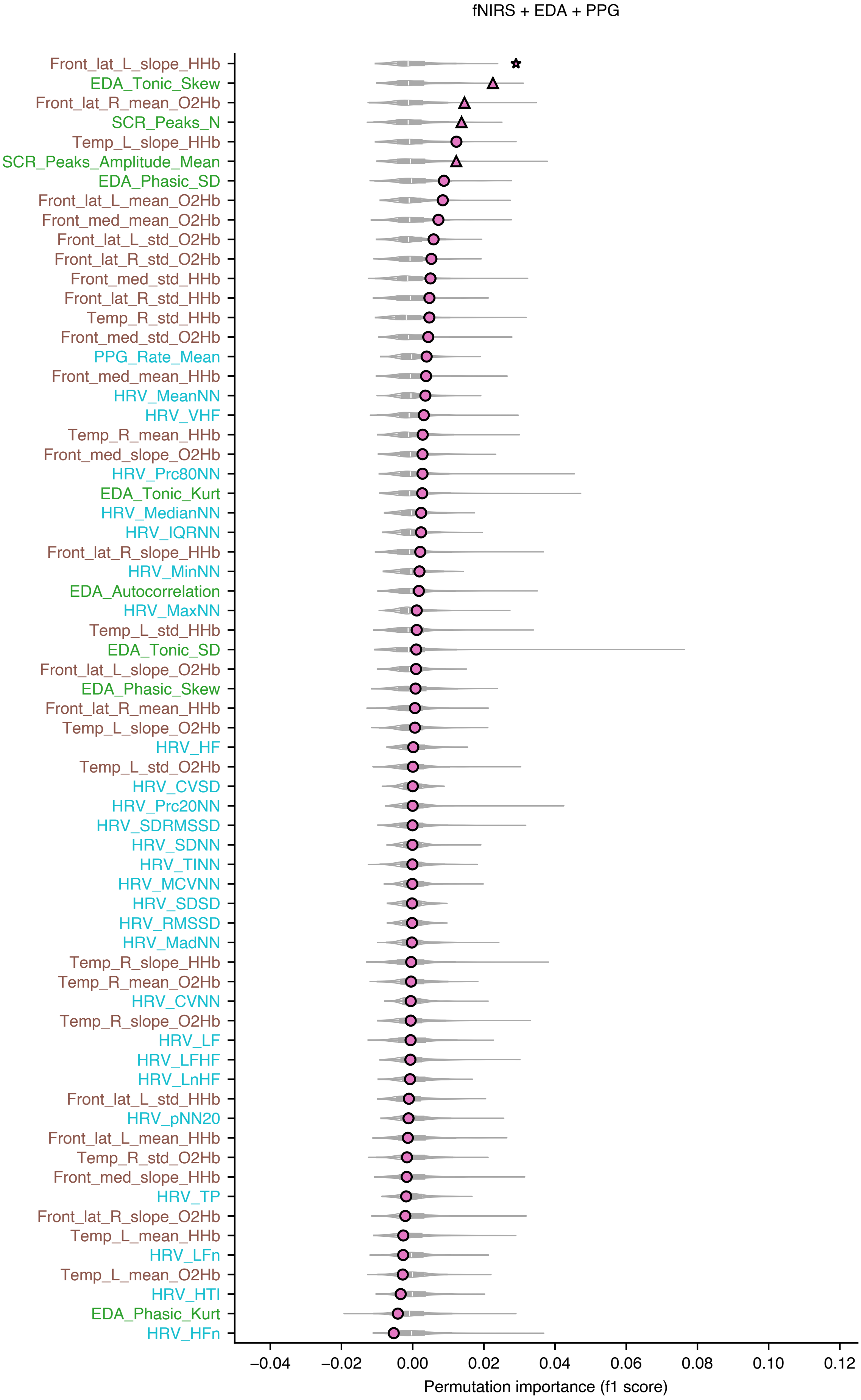

fNIRS + EDA + PPG
Front_lat_L_slope_HHb
EDA_Tonic_Skew
Front_lat_R_mean_O2Hb
SCR_Peaks_N
Temp_L_slope_HHb
SCR_Peaks_Amplitude_Mean
EDA_Phasic_SD
Front_lat_L_mean_O2Hb
Front_med_mean_O2Hb
Front_lat_L_std_O2Hb
Front_lat_R_std_O2Hb
Front_med_std_HHb
Front_lat_R_std_HHb
Temp_R_std_HHb
Front_med_std_O2Hb
PPG_Rate_Mean
Front_med_mean_HHb
HRV_MeanNN
HRV_VHF
Temp_R_mean_HHb
Front_med_slope_O2Hb
HRV_Prc80NN
EDA_Tonic_Kurt
HRV_MedianNN
HRV_IQRNN
Front_lat_R_slope_HHb
HRV_MinNN
EDA_Autocorrelation
HRV_MaxNN
Temp_L_std_HHb
EDA_Tonic_SD
Front_lat_L_slope_O2Hb
EDA_Phasic_Skew
Front_lat_R_mean_HHb
Temp_L_slope_O2Hb
HRV_HF
Temp_L_std_O2Hb
HRV_CVSD
HRV_Prc20NN
HRV_SDRMSSD
HRV_SDNN
HRV_TINN
HRV_MCVNN
HRV_SDSD
HRV_RMSSD
HRV_MadNN
Temp_R_slope_HHb
Temp_R_mean_O2Hb
HRV_CVNN
Temp_R_slope_O2Hb
HRV_LF
HRV_LFHF
HRV_LnHF
Front_lat_L_std_HHb
HRV_pNN20
Front_lat_L_mean_HHb
Temp_R_std_O2Hb
Front_med_slope_HHb
HRV_TP
Front_lat_R_slope_O2Hb
Temp_L_mean_HHb
HRV_LFn
Temp_L_mean_O2Hb
HRV_HTI
EDA_Phasic_Kurt
HRV_HFn
−0.04
−0.02
0.00
0.02
0.04
0.06
0.08
0.10
0.12
Permutation importance (f1 score)

G (Valence)

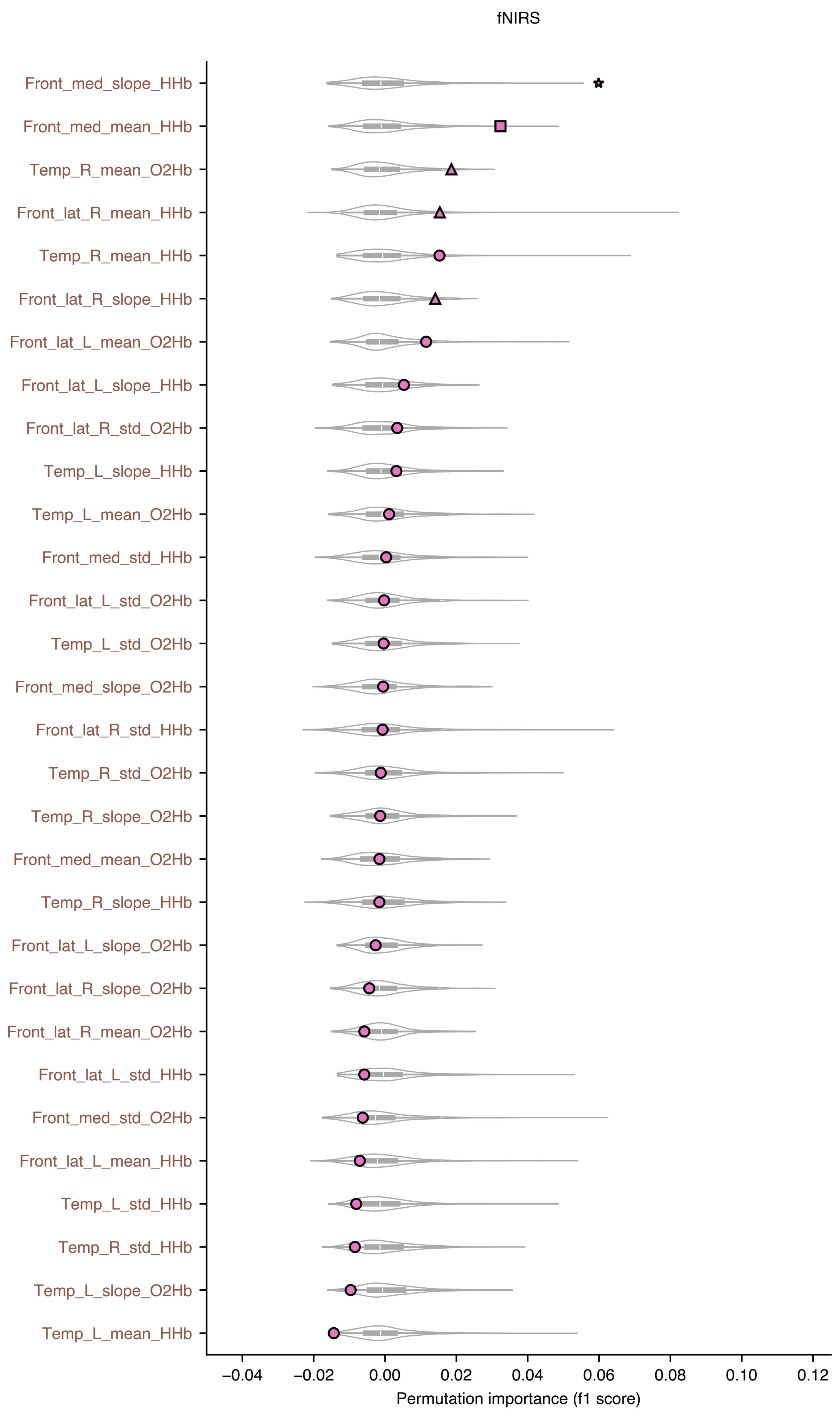

fNIRS
Front_med_slope_HHb
Front_med_mean_HHb
Temp_R_mean_O2Hb
Front_lat_R_mean_HHb
Temp_R_mean_HHb
Front_lat_R_slope_HHb
Front_lat_L_mean_O2Hb
Front_lat_L_slope_HHb
Front_lat_R_std_O2Hb
Temp_L_slope_HHb
Temp_L_mean_O2Hb
Front_med_std_HHb
Front_lat_L_std_O2Hb
Temp_L_std_O2Hb
Front_med_slope_O2Hb
Front_lat_R_std_HHb
Temp_R_std_O2Hb
Temp_R_slope_O2Hb
Front_med_mean_O2Hb
Temp_R_slope_HHb
Front_lat_L_slope_O2Hb
Front_lat_R_slope_O2Hb
Front_lat_R_mean_O2Hb
Front_lat_L_std_HHb
Front_med_std_O2Hb
Front_lat_L_mean_HHb
Temp_L_std_HHb
Temp_R_std_HHb
Temp_L_slope_O2Hb
Temp_L_mean_HHb
−0.04
−0.02
0.00
0.02
0.04
0.06
0.08
0.10
0.12
Permutation importance (f1 score)

H (Valence)

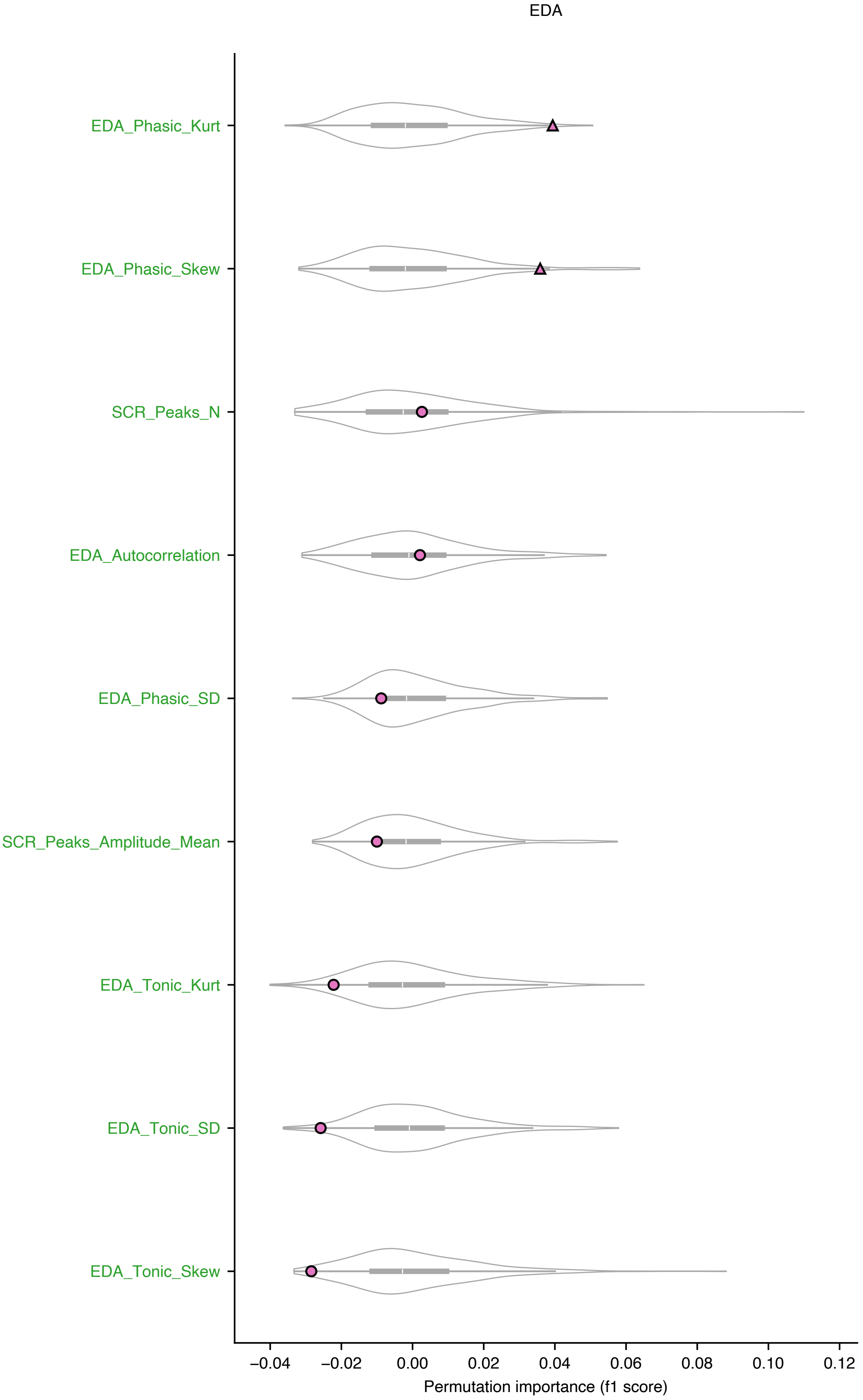
EDA
EDA_Phasic_Kurt
EDA_Phasic_Skew
SCR_Peaks_N
EDA_Autocorrelation
EDA_Phasic_SD
SCR_Peaks_Amplitude_Mean
EDA_Tonic_Kurt
EDA_Tonic_SD
EDA_Tonic_Skew
−0.04
−0.02
0.00
0.02
0.04
0.06
0.08
0.10
0.12
Permutation importance (f1 score)

I (Valence)

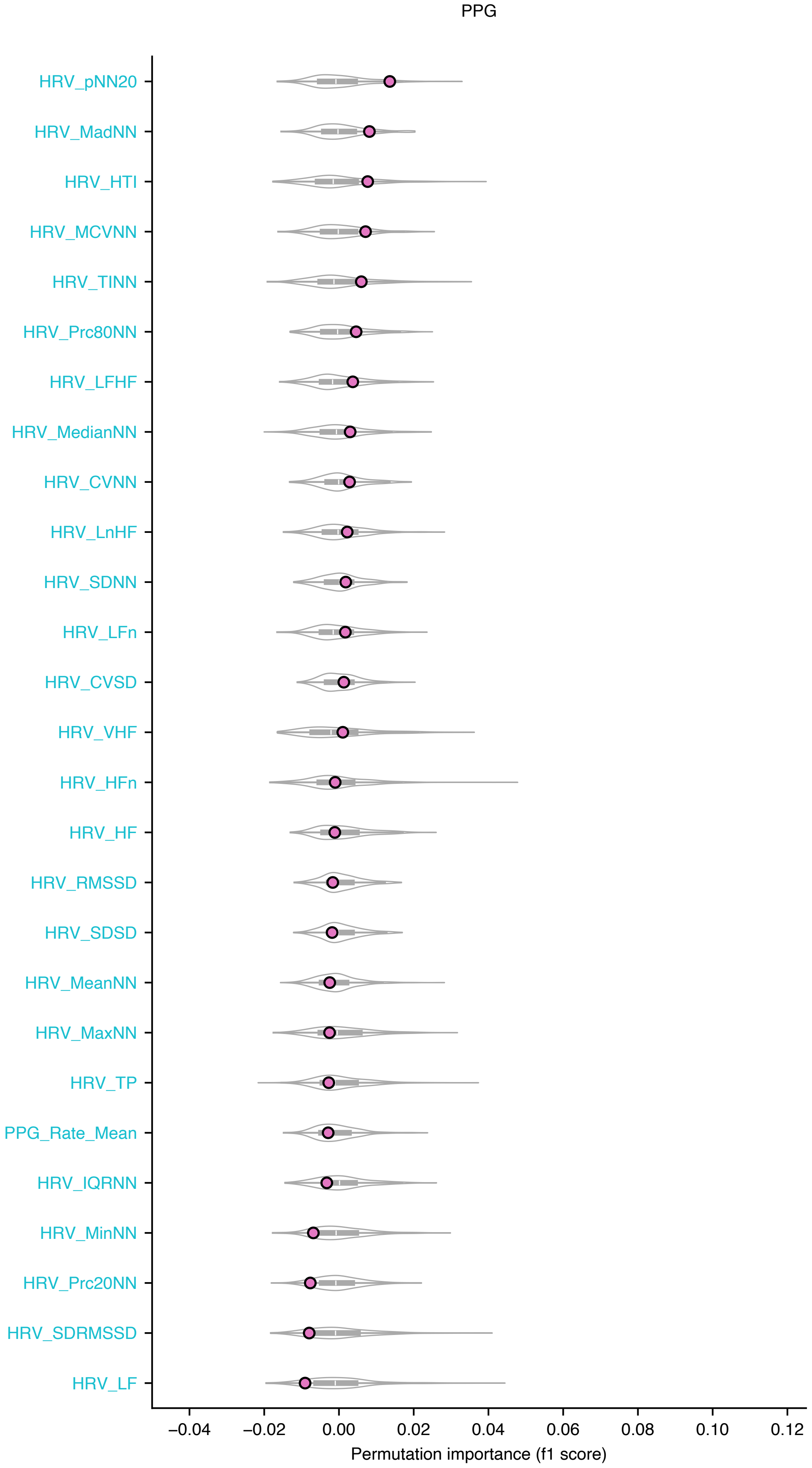

PPG
HRV_pNN20
HRV_MadNN
HRV_HTI
HRV_MCVNN
HRV_TINN
HRV_Prc80NN
HRV_LFHF
HRV_MedianNN
HRV_CVNN
HRV_LnHF
HRV_SDNN
HRV_LFn
HRV_CVSD
HRV_VHF
HRV_HFn
HRV_HF
HRV_RMSSD
HRV_SDSD
HRV_MeanNN
HRV_MaxNN
HRV_TP
PPG_Rate_Mean
HRV_IQRNN
HRV_MinNN
HRV_Prc20NN
HRV_SDRMSSD
HRV_LF
−0.04
−0.02
0.00
0.02
0.04
0.06
0.08
0.10
0.12
Permutation importance (f1 score)

J (Valence)

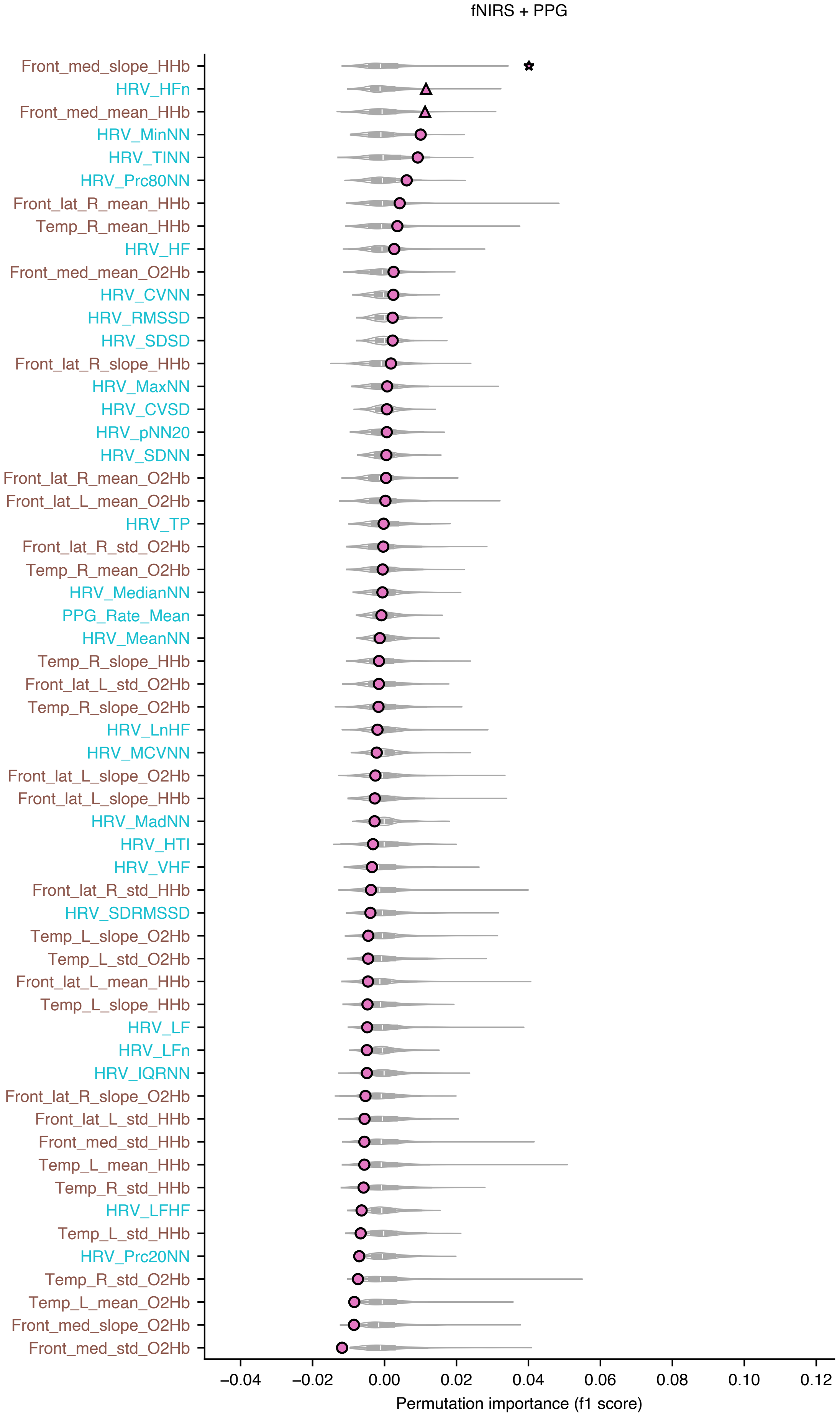

fNIRS + PPG
Front_med_slope_HHb
HRV_HFn
Front_med_mean_HHb
HRV_MinNN
HRV_TINN
HRV_Prc80NN
Front_lat_R_mean_HHb
Temp_R_mean_HHb
HRV_HF
Front_med_mean_O2Hb
HRV_CVNN
HRV_RMSSD
HRV_SDSD
Front_lat_R_slope_HHb
HRV_MaxNN
HRV_CVSD
HRV_pNN20
HRV_SDNN
Front_lat_R_mean_O2Hb
Front_lat_L_mean_O2Hb
HRV_TP
Front_lat_R_std_O2Hb
Temp_R_mean_O2Hb
HRV_MedianNN
PPG_Rate_Mean
HRV_MeanNN
Temp_R_slope_HHb
Front_lat_L_std_O2Hb
Temp_R_slope_O2Hb
HRV_LnHF
HRV_MCVNN
Front_lat_L_slope_O2Hb
Front_lat_L_slope_HHb
HRV_MadNN
HRV_HTI
HRV_VHF
Front_lat_R_std_HHb
HRV_SDRMSSD
Temp_L_slope_O2Hb
Temp_L_std_O2Hb
Front_lat_L_mean_HHb
Temp_L_slope_HHb
HRV_LF
HRV_LFn
HRV_IQRNN
Front_lat_R_slope_O2Hb
Front_lat_L_std_HHb
Front_med_std_HHb
Temp_L_mean_HHb
Temp_R_std_HHb
HRV_LFHF
Temp_L_std_HHb
HRV_Prc20NN
Temp_R_std_O2Hb
Temp_L_mean_O2Hb
Front_med_slope_O2Hb
Front_med_std_O2Hb
−0.04
−0.02
0.00
0.02
0.04
0.06
0.08
0.10
0.12
Permutation importance (f1 score)

K (Valence)

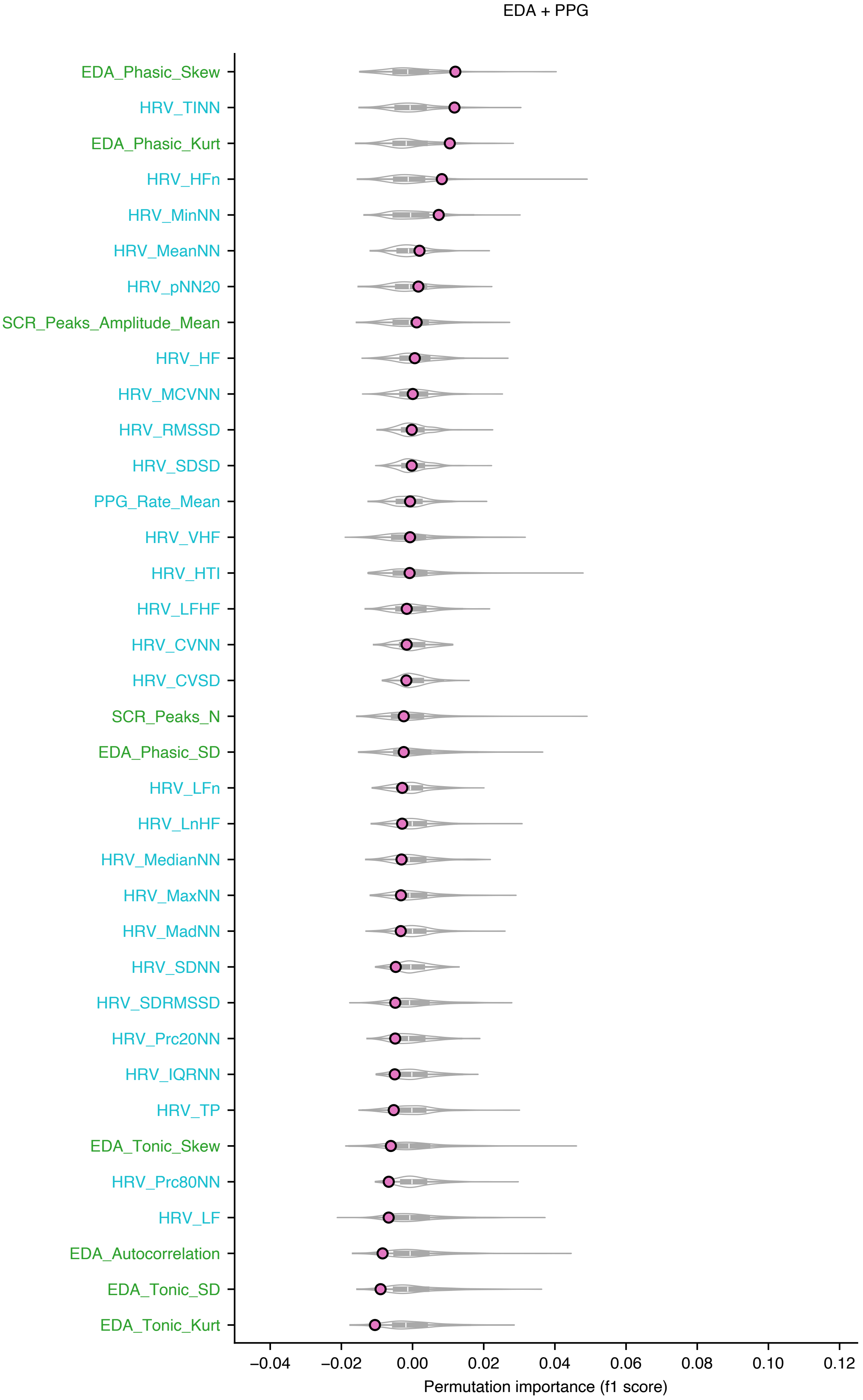
EDA + PPG
EDA_Phasic_Skew
HRV_TINN
EDA_Phasic_Kurt
HRV_HFn
HRV_MinNN
HRV_MeanNN
HRV_pNN20
SCR_Peaks_Amplitude_Mean
HRV_HF
HRV_MCVNN
HRV_RMSSD
HRV_SDSD
PPG_Rate_Mean
HRV_VHF
HRV_HTI
HRV_LFHF
HRV_CVNN
HRV_CVSD
SCR_Peaks_N
EDA_Phasic_SD
HRV_LFn
HRV_LnHF
HRV_MedianNN
HRV_MaxNN
HRV_MadNN
HRV_SDNN
HRV_SDRMSSD
HRV_Prc20NN
HRV_IQRNN
HRV_TP
EDA_Tonic_Skew
HRV_Prc80NN
HRV_LF
EDA_Autocorrelation
EDA_Tonic_SD
EDA_Tonic_Kurt
−0.04
−0.02
0.00
0.02
0.04
0.06
0.08
0.10
0.12
Permutation importance (f1 score)

L (Valence)

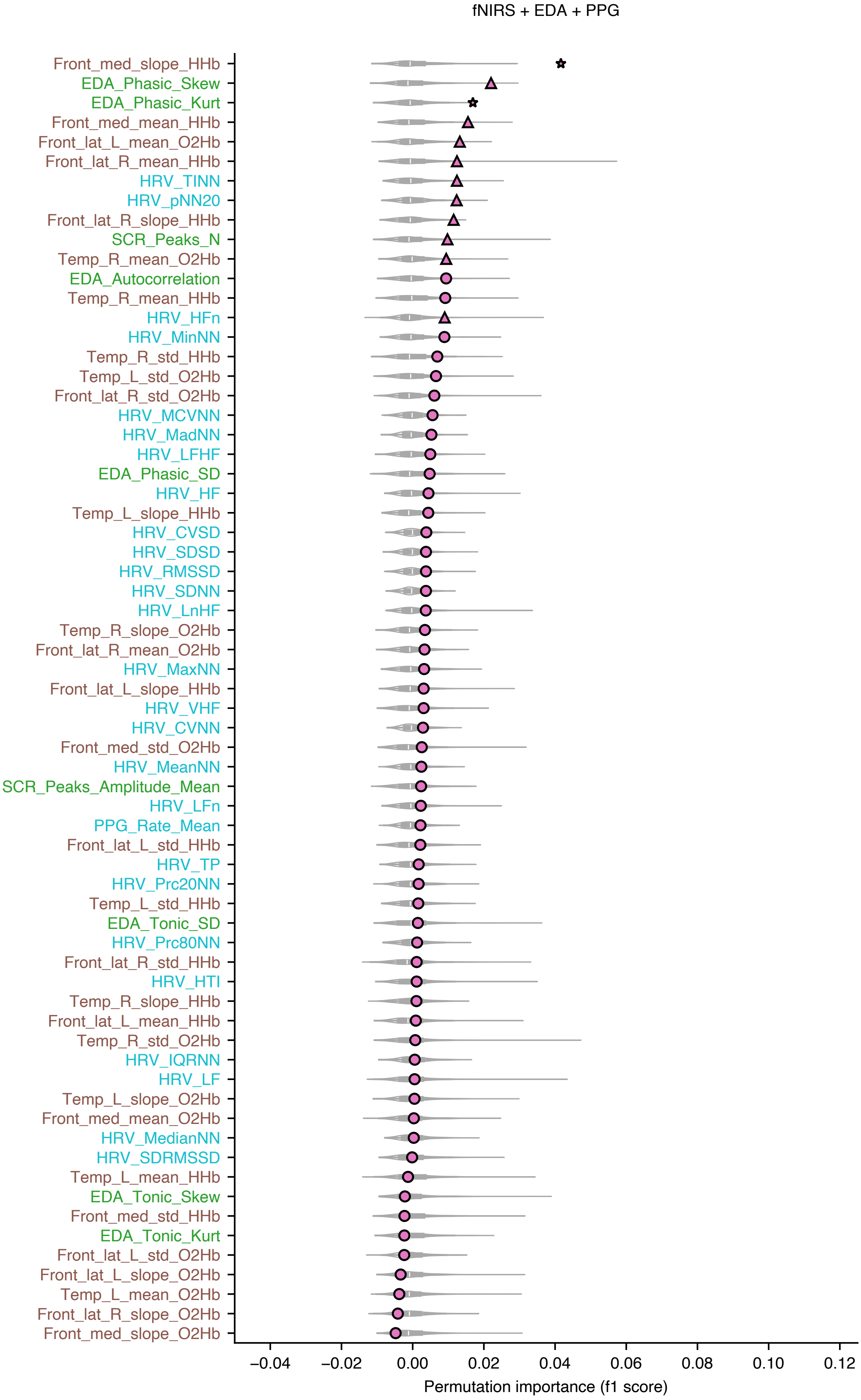
fNIRS + EDA + PPG
Front_med_slope_HHb
EDA_Phasic_Skew
EDA_Phasic_Kurt
Front_med_mean_HHb
Front_lat_L_mean_O2Hb
Front_lat_R_mean_HHb
HRV_TINN
HRV_pNN20
Front_lat_R_slope_HHb
SCR_Peaks_N
Temp_R_mean_O2Hb
EDA_Autocorrelation
Temp_R_mean_HHb
HRV_HFn
HRV_MinNN
Temp_R_std_HHb
Temp_L_std_O2Hb
Front_lat_R_std_O2Hb
HRV_MCVNN
HRV_MadNN
HRV_LFHF
EDA_Phasic_SD
HRV_HF
Temp_L_slope_HHb
HRV_CVSD
HRV_SDSD
HRV_RMSSD
HRV_SDNN
HRV_LnHF
Temp_R_slope_O2Hb
Front_lat_R_mean_O2Hb
HRV_MaxNN
Front_lat_L_slope_HHb
HRV_VHF
HRV_CVNN
Front_med_std_O2Hb
HRV_MeanNN
SCR_Peaks_Amplitude_Mean
HRV_LFn
PPG_Rate_Mean
Front_lat_L_std_HHb
HRV_TP
HRV_Prc20NN
Temp_L_std_HHb
EDA_Tonic_SD
HRV_Prc80NN
Front_lat_R_std_HHb
HRV_HTI
Temp_R_slope_HHb
Front_lat_L_mean_HHb
Temp_R_std_O2Hb
HRV_IQRNN
HRV_LF
Temp_L_slope_O2Hb
Front_med_mean_O2Hb
HRV_MedianNN
HRV_SDRMSSD
Temp_L_mean_HHb
EDA_Tonic_Skew
Front_med_std_HHb
EDA_Tonic_Kurt
Front_lat_L_std_O2Hb
Front_lat_L_slope_O2Hb
Temp_L_mean_O2Hb
Front_lat_R_slope_O2Hb
Front_med_slope_O2Hb
−0.04
−0.02
0.00
0.02
0.04
0.06
0.08
0.10
0.12
Permutation importance (f1 score)
